# Cdc42 small GTPase is a novel regulator of the fibrogenic activation of human intestinal myofibroblasts

**DOI:** 10.64898/2026.07.09.737543

**Authors:** Atif Zafar, Gaurav Chauhan, Pranab K. Mukherjee, Armando Marino-Melendez, Ryan Musich, Yan Wang, Nayden G. Naydenov, Florian Rieder, Andrei I. Ivanov

## Abstract

Cell division cycle 42 (Cdc42) is a member of the Rho family of small GTPases, which plays crucial roles in regulating cytoskeletal remodeling, and membrane trafficking. While previous studies implicated Cdc42 in controlling intestinal epithelial homeostasis, the involvement of this small GTPase in the process of intestinal fibrogenesis remains unexplored. Our study was designed to determine whether Cdc42 regulates the fibrogenic activation of intestinal myofibroblasts *in vitro*. The study was conducted using a CCD-18Co normal human colonic fibroblast cell line, and primary human intestinal myofibroblasts (HIMF) isolated from Crohn’s disease (CD) patients. CCD-18Co and HIMF cells were stimulated by transforming growth factor-β1 (TGF-β1). Cdc42 was inhibited either genetically, using siRNA-mediated knockdown, or pharmacologically using specific Cdc42 inhibitors, ML141 and CASIN. Genetic and pharmacologic inhibition of Cdc42 markedly reduced TGF-β1 induced expression of the major contractile cytoskeletal proteins, α-smooth muscle actin, calponin 1 and L-caldesmon. Furthermore, Cdc42 inhibition significantly attenuated expression of key extracellular matrix (ECM) proteins, fibronectin and collagen I, in activated CCD-18Co cells and HIMF. Interestingly, decreased expression of contractile and ECM proteins in Cdc42-depleted myofibroblasts was not due to downregulation of the TGF-β1 signaling, decreased mRNA transcription or increased lysosomal or proteasomal degradation of these proteins. Such suppressed pro-fibrotic activation of Cdc42-deficient CCD-18Co cells and HIMF involved a selective inhibition of protein translation due to inactivation of the AKT-mammalian target of rapamycin (mTOR) signaling module. These findings highlight Cdc42 as a key regulator of intestinal fibrosis that controls mTOR activation to enhance ECM production and contractile actomyosin cytoskeleton in intestinal myofibroblasts.

**In brief:** Zafar et al. unravel a novel role of Cdc42 small GTPase in fibrogenic activation of human intestinal myofibroblasts. Genetic and pharmacologic inhibition of Cdc42 markedly reduced TGF-β induced expression of contractile cytoskeletal proteins and extracellular matrix proteins in activated myofibroblasts. The mechanisms underlying such profibrotic activity of Cdc42 involve regulation of de novo protein translation via activation of the AKT-mTOR signaling module.

**Highlights:**

- Cdc42 regulates TGF-β dependent activation of human intestinal myofibroblasts
- Profibrotic activity of Cdc42 involves regulation of de novo protein translation
- Cdc42 regulates myofibroblast activation via AKT–mTOR signaling pathway
- Targeting Cdc42–Akt–mTOR signaling may facilitate the development of novel antifibrotic therapies

## Introduction

Intestinal fibrosis is a major complication of inflammatory bowel diseases (IBD), in particular in Crohn’s disease (CD)^1,2^. Fibrosis builds up in the intestine results in stiffening of the gut wall and narrowing the gut lumen, thereby creating intestinal strictures that obstructs luminal transit leading to surgery. Currently, there are no drugs to selectively target intestinal fibrosis^3,4^. Thus, a better understanding of the molecular mechanisms of intestinal fibrosis is of paramount importance for the development of future treatment strategies. Fibrosis of the intestine and other organs develops due to excessive production of extracellular matrix (ECM), containing collagen I, fibronectin and other secreted proteins. Myofibroblasts are considered major drivers of the intestinal fibrosis^5,6^. This cell type originates via activation of fibroblasts and other intestinal stromal cells by tissue growth factors, inflammatory mediators, as well as luminal bacterial products^7,8^. Transforming growth factor-beta (TGF-β) is a *bona fide* activator of myofibroblasts and a major pro-fibrotic molecule in the intestine and other organs^9,10^.

TGF-β induced myofibroblast activation involves increased production of ECM proteins, such as fibronectin and collagen I^11–13^, along with the acquisition of a contractile, smooth muscle cell-like phenotype^11–13^. These unique functional features of activated myofibroblasts are driven by the fundamental cellular mechanisms such as increased vesicle trafficking and secretion and enhancement/remodeling of the actomyosin cytoskeleton^14–16^. Understanding the molecular/signaling pathways that are responsible for such trafficking and cytoskeletal enhancement is essential to prevent and/or reverse myofibroblast activation, potentially leading to novel antifibrotic therapies.

Rho family of small GTPases serves as a critical signaling hub receiving imputes from various extracellular stimuli to activate the intracellular signaling pathways. The Rho-activated signaling pathways control virtually all major cellular processes including cytoskeletal assembly/remodeling and membrane trafficking^17–19^. RhoA, Rac1 and Cdc42 are the founding and the most studied members of the Rho family of small GTPases. Among them, RhoA and Rac1 are well known regulators of myofibroblast activation and fibrosis of the intestine and other organs^20–27^. By contrast, the fibrogenic roles of Cdc42 remain largely understudied, representing a major knowledge gap in the field.

Similarly to other Rho small GTPases, Cdc42 acts as a molecular switch, being active in a GTP-bound form and inactivated after the GTP to GDP hydrolysis^28–30^. Cdc42 readily binds to phospholipid membranes and is commonly activated at the cell cortex by plasma membrane receptor tyrosine kinases, G-protein coupled receptors and integrins^28–30^. Cdc42 is known to regulate a variety of important cellular processes such as cell division, migration, cell-cell adhesions and cell polarity by activating several downstream effector molecules^28–30^. Importantly Cdc42 is a key regulator of the actomyosin cytoskeleton targeting both actin filament assembly and contraction^28–30^. Furthermore, this Rho GTPase was shown to control intracellular vesicle trafficking and protein secretion^17,31^. Such dual control of the cytoskeleton and vesicle trafficking makes Cdc42 a potentially important regulator of myofibroblast activation; however, its role has only recently begun to emerge and remains poorly understood. No previous studies addressed Cdc42 function in intestinal fibrogenesis, while recent publications suggest either inhibition^32–34^ or acceleration^35,36^ of renal and liver fibrosis by activated Cdc42.

Our study has been designed to fill this important knowledge gap and investigate the roles of Cdc42 in TGF-β dependent activation of human intestinal myofibroblasts (HIMF). We report that Cdc42 activation determines two major molecular signatures of HIMF activation: (1) increased production/deposition of ECM and (2) enhanced expression of contractile cytoskeletal proteins. The mechanisms underlying such profibrotic activity of Cdc42 involve regulation of de novo protein translation via activation of the AKT-mTOR signaling module.

## Results

### Cdc42 is abundantly expressed in distinct stromal cell populations in the human intestine

Tissue fibrosis is known to be driven by activation of different stromal cell populations. In order to characterize Cdc42 in cell types relevant to intestinal fibrosis, we profiled expression of this small GTPase in intestinal stromal cells using our recent single-cell RNA sequencing (scRNA-seq) data^37^. Analysis of full-thickness sections of normal human colon revealed Cdc42 expression in different stromal cell types, being abundant in vascular and lymphatic endothelium, fibroblasts and smooth muscle cells (Figure 1A). We next asked if Cdc42 expression is altered in different intestinal stromal cell populations of patients with Crohn’s disease. We separately evaluated non-inflamed, inflamed but non-strictured and strictured Crohn’s disease tissues (designated as CDni, CDi and CDs, respectively) as compared to non-IBD control tissues (designated as NL). Several populations of CDs fibroblasts demonstrated increased Cdc42 expression compared to normal controls (Figure 1B). Next, we focused on the effects of fibrotic milieu of intestinal stroma on Cdc42 expression and identified two populations of fibroblasts (fibroblast-ECM and fibroblast-reticular cells) and a population of endothelial cells (endothelial-venular) with high Cdc42 expression in fibrotic CDs as compared to non-fibrotic CDni tissue (Suppl. Figure 1). Pathway analysis in these stromal cell populations revealed upregulation of Cdc42 activation pathways along with several downstream Cdc42 effectors (IQGAPs, PAKs and N-WASP/WAVE) in strictured CD fibroblasts (Suppl. Figure 2). Interestingly, these effects appear to be fibroblast-specific and were not recapitulated in the Cdc42-high endothelial cell population (Suppl. Figure 2). Overall, these data suggest that Cdc42 is expressed in a variety of human intestinal stromal cells and that its signaling could be enhanced in subpopulations of fibroblasts in fibrotic CD strictures.

**Figure 1.**
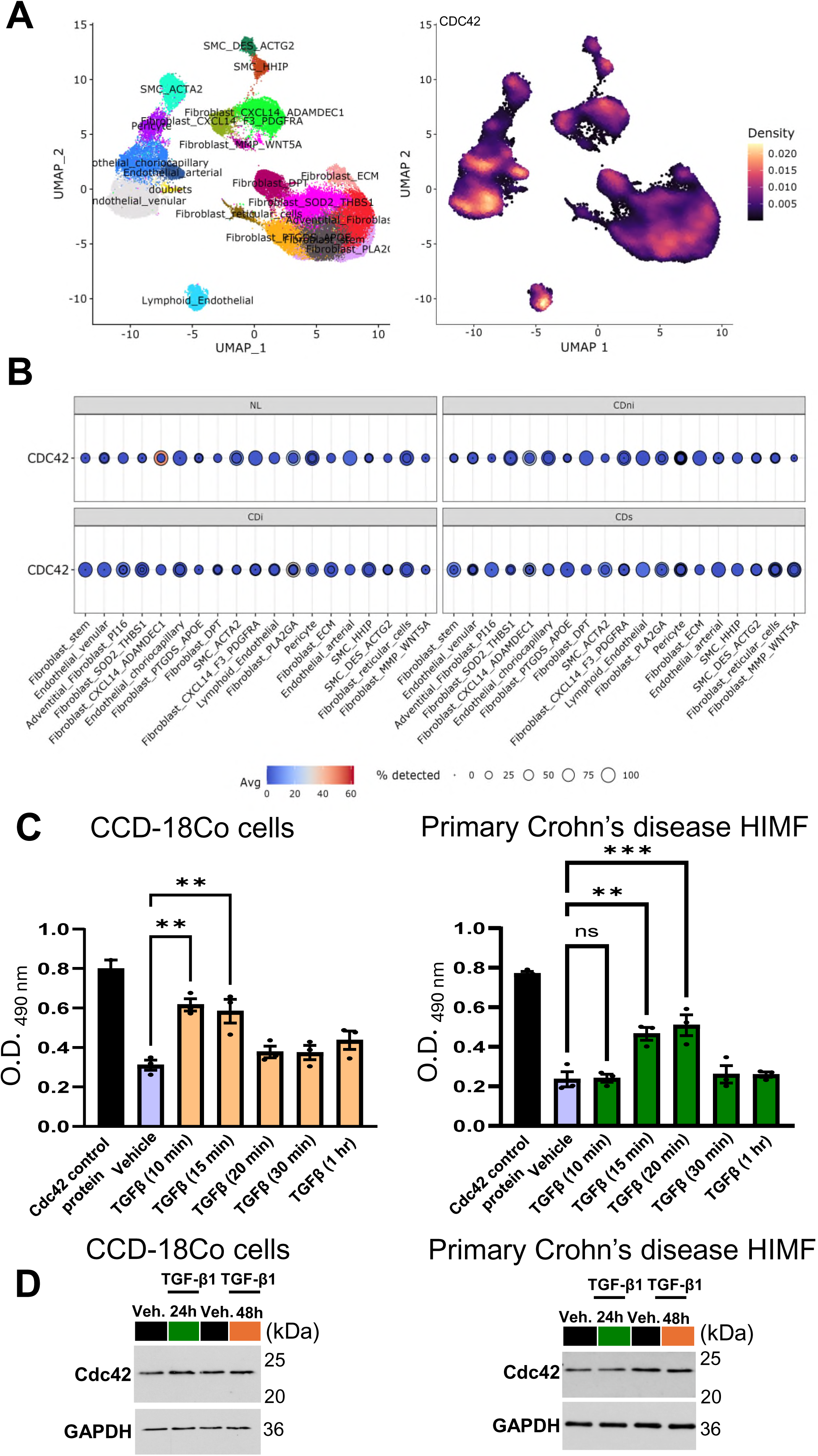
Cdc42 is abundantly expressed in colonic stromal cells and is activated by TGF-β in human intestinal myofibroblasts. (**A**) Uniform manifold approximation and projection (UMAP) plot (left) and density plot (right) representation of Cdc42 expression in stromal cells in normal human colonic mucosa. (**B**) Dot plot showing increased expression of Cdc42 in fibroblast subpopulation in strictured Crohn’s disease (CDs). NL: non-IBD patients; CDni: nonstrictured, noninflamed Crohn’s disease; CDi: nonstrictured, inflamed Crohn’s disease; CDs strictured Crohn’s disease. (**C and D**) CCD-18Co cells or primary HIMF isolated from CD patients were stimulated by TGF-β1 for indicated times. The amount of active Cdc42 was determined by G-ELISA assay **(C)**. The effect of long term TGF-β1 stimulation on total Cdc42 expression was determine by immunoblotting (**D**). Values are reported as mean ± standard error of mean (SEM) (n=3). *p < 0.05, **p < 0.01 and ns, not significant by one-way ANOVA with Tukey’s multiple-comparison test.

### Cdc42 is activated by TGF-β in cultured colonic fibroblasts

Next, we determined whether Cdc42 is activated by profibrotic TGF-β signaling in human intestinal fibroblasts using two different cell types. One is CCD-18Co cells, which are normal human colonic fibroblasts commonly used to examine molecular and cellular mechanisms of intestinal fibrosis^38–42^. The second cell type is primary human intestinal myofibroblasts (HIMF) isolated from resected intestinal tissues of patients with inflamed and strictured CD. Treatment of CCD18-Co and HIMF cells with TGF-β1 caused a rapid transient activation of Cdc42 according to G-ELISA GTPase assays (Figure 1C). The amount of active Cdc42 peaked at 10-20 min of the treatment rapidly decreasing thereafter. No significant changes in expression of total Cdc42 protein was detected even after 24 or 48 h of TGF-β1 treatment (Figure 1D). These data suggest that Cdc42 activation represents an early event of TGF-β dependent myofibroblast activation.

### Pharmacologic or genetic inhibition of Cdc42 blocks the fibrogenic activation of intestinal fibroblasts

A combination of pharmacological inhibition and genetic depletion of Cdc42 was used to determine whether this small GTPase is essential for TGF-β induced fibrogenic activation of intestinal myofibroblasts. CCD-18Co cells and HIMF were exposed to TGF-β1 for 48 h in the presence of either selective Cdc42 inhibitors ML141 or casin^43–47^, or vehicle. Fibroblast activation was determined by examining cellular levels of contractile cytoskeletal proteins, α-SMA, calponin-1 and L-caldesmon as well as expression of key ECM proteins, collagen I and fibronectin (Figure 2A and B). In CCD-18Co cells, both Cdc42 inhibitors significantly attenuated TGF-β1 induced expression of all tested contractile myofibroblast markers and ECM proteins (Figure 2A and B). Similar inhibitory effects were observed in activated HIMF on both contractile myofibroblast markers and ECM proteins following exposure with the Cdc42 inhibitors ML141 or casin (Figure 2A and B). As a complimentary approach we used an ECM deposition assay to examine if Cdc42 inhibition affects secretion of collagen I and fibronectin. TGF-β1 treatment of primary HIMF from CD patients caused robust deposition of collagen I and fibronectin and such matrix deposition was markedly attenuated by both ML141 and casin (Figure 2C).

**Figure 2.**
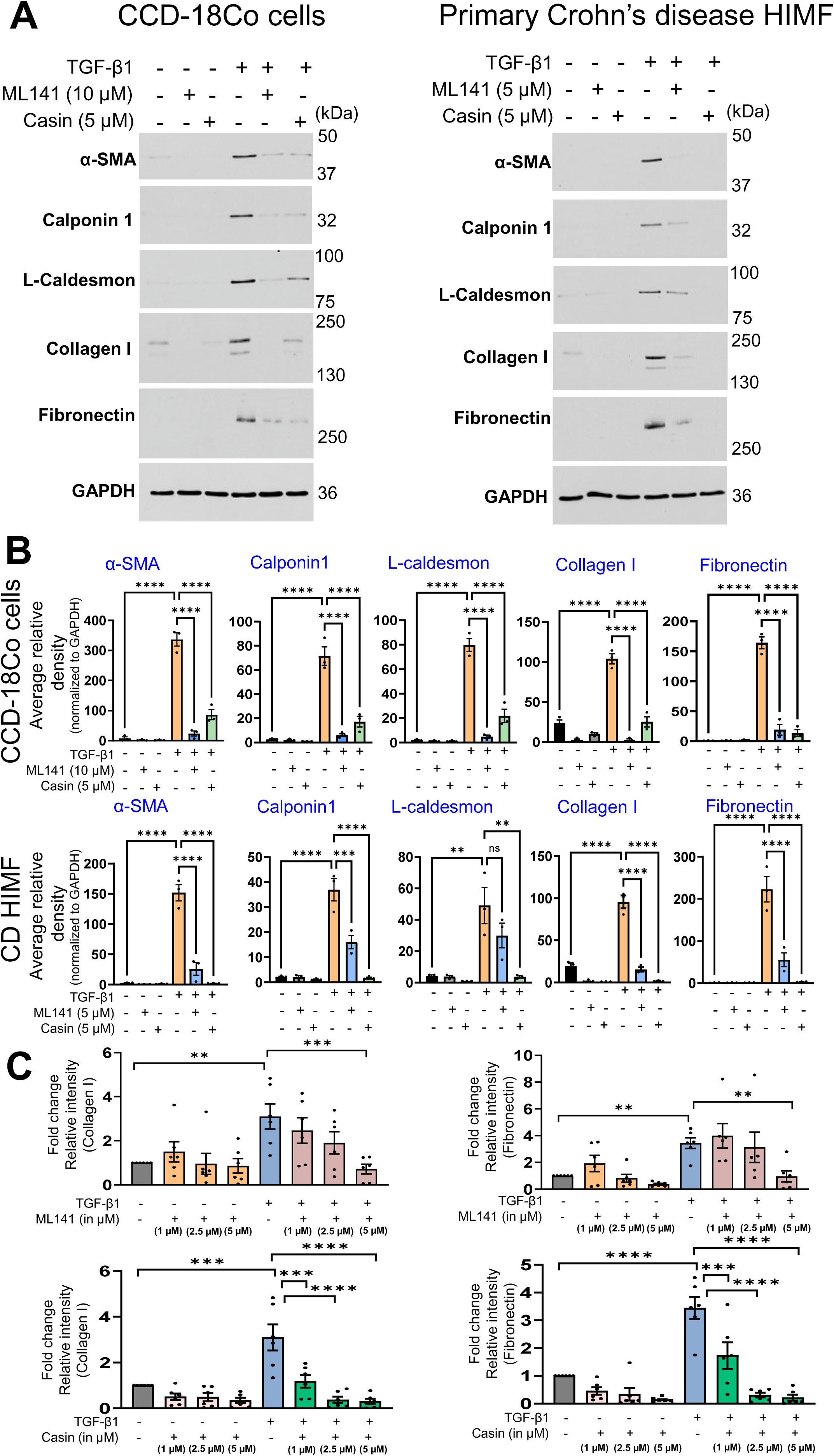
Pharmacological inhibition of Cdc42 attenuates TGF-β1-induced activation of human intestinal myofibroblasts and their extracellular matrix (ECM) secretion. CCD-18Co cells and primary CD HIMF were treated for 48 hours with TGF-β1 in the presence of either pharmacologic Cdc42 inhibitors ML141 and casin or vehicle. (**A**) Representative immunoblotting images and **(B)** densitometric quantification of protein expression of cytoskeletal myofibroblast markers (α-SMA, calponin 1, L-caldesmon) and ECM proteins (collagen I and fibronectin) in total cell lysates. Average relative band densities represent the values normalized to corresponding GAPDH relative band densities (loading control) (n=3). (**C**) ECM deposition assay of primary CD HIMF activated in the presence of different concentrations of Cdc42 inhibitors (ML141 or casin) (n=6). Data in **(B)** were analyzed by one-way ANOVA, and data in **(C)** was analyzed by two-way ANOVA followed by Tukey’s multiple-comparison test. All data are expressed as mean ± SEM. ^∗^p < 0.05, ^∗∗^p < 0.01, ^∗∗∗^p < 0.001.

To verify our pharmacologic inhibition data with more selective genetic inhibition of Cdc42, we downregulated the expression of this small GTPase by using RNA interference. Immunoblotting analysis revealed that Cdc42 depletion significantly attenuated TGF-β1 induced expression of contractile markers and ECM proteins in CCD-18Co cells (Figure 3A and B) and primary HIMF (Figure 3A and B). Inhibition of TGF-β1 induced expression of α-SMA and fibronectin in Cdc42-depleted HIMF was also observed by immunofluorescence labeling and confocal microscopy (Suppl. Figure 3). Finally, the ECM deposition assay revealed a substantial inhibition of collagen I and fibronectin secretion by TGF-β1 stimulated HIMF following Cdc42 depletion (Figure 3C). Importantly, the observed inhibitory effects of either pharmacological inhibition or genetic depletion were not attributable to off-target cellular cytotoxicity, as demonstrated by LDH assay and western blot analysis of apoptosis markers (Suppl. Figure 4). Together, pharmacologic and genetic inhibition data demonstrate an important role of Cdc42 as a mediator of TGF-β-induced fibrogenic activation of human intestinal myofibroblasts.

**Figure 3.**
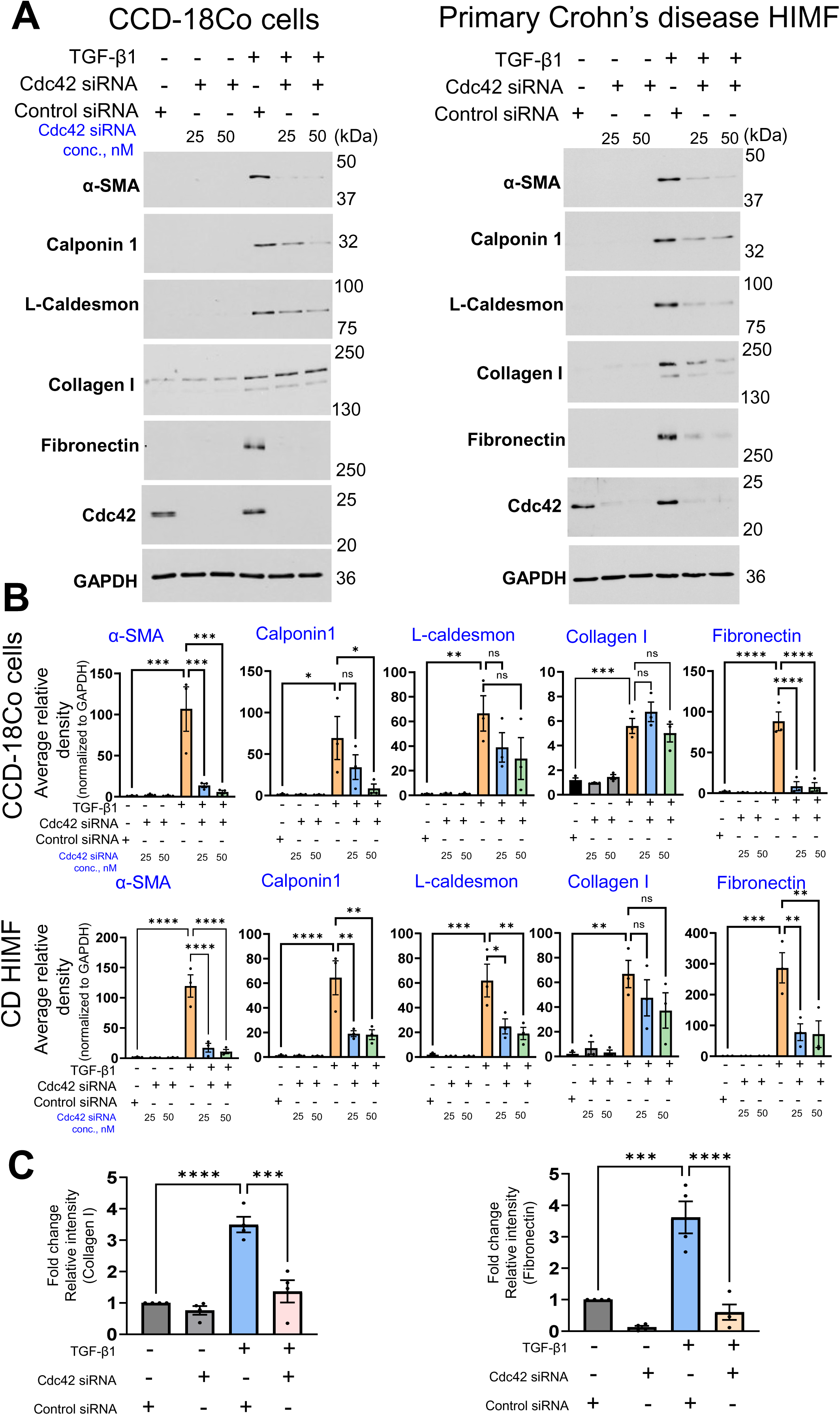
Genetic downregulation of Cdc42 expression attenuates TGF-β1-induced activation of human intestinal myofibroblasts. Control and Cdc42 depleted CCD-18Co cells and primary CD HIMF were activated for 48 hours with TGF-β1. (**A and B**) Representative immunoblotting images and **(B)** densitometric quantification of protein expression of cytoskeletal myofibroblast markers (α-SMA, calponin 1, L-caldesmon) and extracellular matrix proteins (collagen I and fibronectin) in total cell lysates. Average relative band densities represent the values normalized to corresponding GAPDH relative band densities (loading control) (n=3). (**C**) ECM deposition assay in control and Cdc42-depleted primary CD HIMF under resting conditions and after TGF-β activation (n=4). Data are presented as mean ± SEM. *p < 0.05; **p < 0.01; ***p < 0.001 by one-way ANOVA with Tukey’s multiple-comparison test.

### Cdc42 depletion does not inhibit TGF-β induced signaling, and has no effects on transcription or degradation of contractile and ECM proteins

Next, we sought to examine the mechanisms underlying Cdc42-dependent regulation of intestinal myofibroblast activation. Since increase in Cdc42 activity is an early molecular event in TGF-β stimulated fibroblasts, we asked whether it could affect signaling via TGF-β receptor by examining SMAD2/3 phosphorylation. In both CCD-18Co cells and primary CD HIMF, TGF-β1 triggered a rapid and transient increase in SMAD2/3 phosphorylation that peaked at 30-60 min of the exposure (Suppl. Figure 5). Cdc42 depletion affected neither the total levels of phospho-SMAD2/3 proteins (Figure 4A), nor the translocation of activated SMAD2 into the nucleus in CCD-18Co cells and primary CD HIMF (Figure 4B). Consistently, loss of Cdc42 did not attenuate TGF-β1 induced mRNA transcription of α-SMA, fibronectin and collagen I (Figure 4C). Together, these data strongly suggest that Cdc42 does not control TGF-β signaling and transcription of major TGF-β induced genes in activated intestinal myofibroblasts.

**Figure 4.**
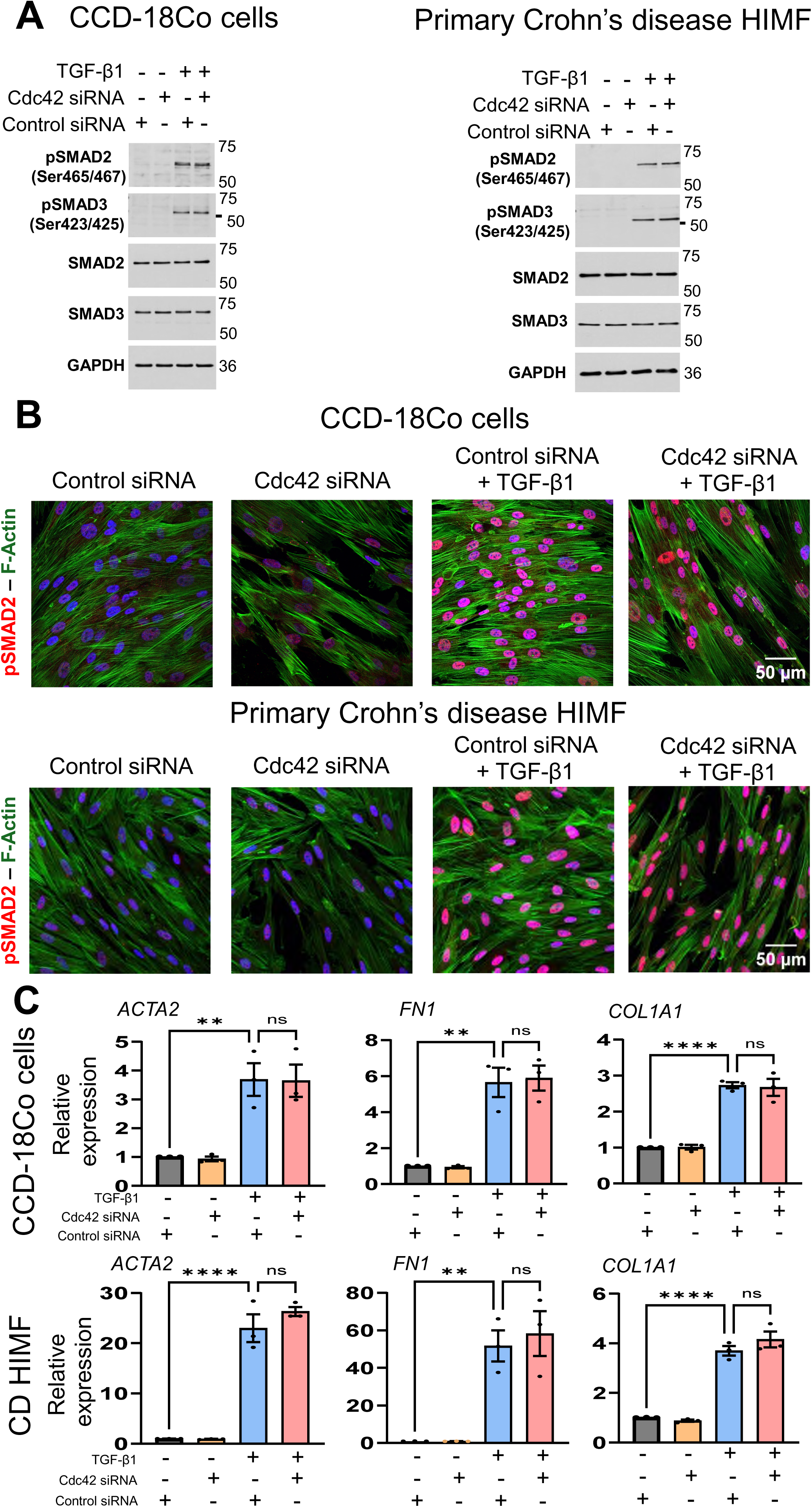
Depletion of Cdc42 signaling does not affect SMAD activation and transcription of contractile and ECM proteins in TGF-β stimulated intestinal myofibroblasts. (**A**) Control and Cdc42-depleted CCD-18Co cells and primary CD HIMF were stimulated with TGF-β1 for 1 hour and expression of total and phosphorylated SMAD2 and SMAD3 proteins was determined by immunoblotting analysis. (**B**) Effect of TGF-β1 on the nuclear translocation of pSMAD2 was determined by fluorescence labeling for pSMAD2 (red), F-actin (green) and nuclei (blue) in control or Cdc42-depleted intestinal myofibroblasts. (**C**) CCD-18Co cells and CD HIMF were transfected with control or Cdc42-specific siRNAs and stimulated with TGF-β1 for 48 hours. Total RNA was isolated and analyzed for the expression *ACTA2, FN1 and COL1A1* by quantitative qRT-PCR. Data are presented as mean ± SEM (n=3). *p < 0.05; **p < 0.01; ***p < 0.001 by one-way ANOVA with Tukey’s multiple-comparison test. Scale bars, 50 µm.

Another mechanism that could underline Cdc42 functions in TGF-β stimulated intestinal myofibroblasts involves regulation of protein degradation. We tested this mechanism by using selective inhibitors of two major degradation pathways, namely, lysosomal and proteasomal degradation. Lysosomal degradation was inhibited by bafilomycin A1 treatment. The selected bafilomycin dose and treatment duration efficiently blocked lysosomal functions based on the increased levels of the autophagosomal markers, lipidated LC3 and p62 protein (Figure 5A). However, such inhibition of lysosomal degradation failed to restore the decrease in expression of α-SMA, collagen I and fibronectin in activated CCD-18Co cells and primary HIMF caused by Cdc42 depletion (Figure 5A and B). Likewise, a proteasomal degradation inhibitor, MG132, efficiently blocked proteasomal functions based on upregulated expression of different ubiquitinated proteins (Figure 5C). However, this inhibitor also failed to restore α-SMA, collagen I and fibronectin expression in Cdc42 depleted cells exposed to TGF-β1 (Figure 5C and D). Together these data argue against the idea that Cdc42 inhibition downregulates expression of contractile and ECM proteins by stimulating their degradation in activated intestinal myofibroblasts.

**Figure 5.**
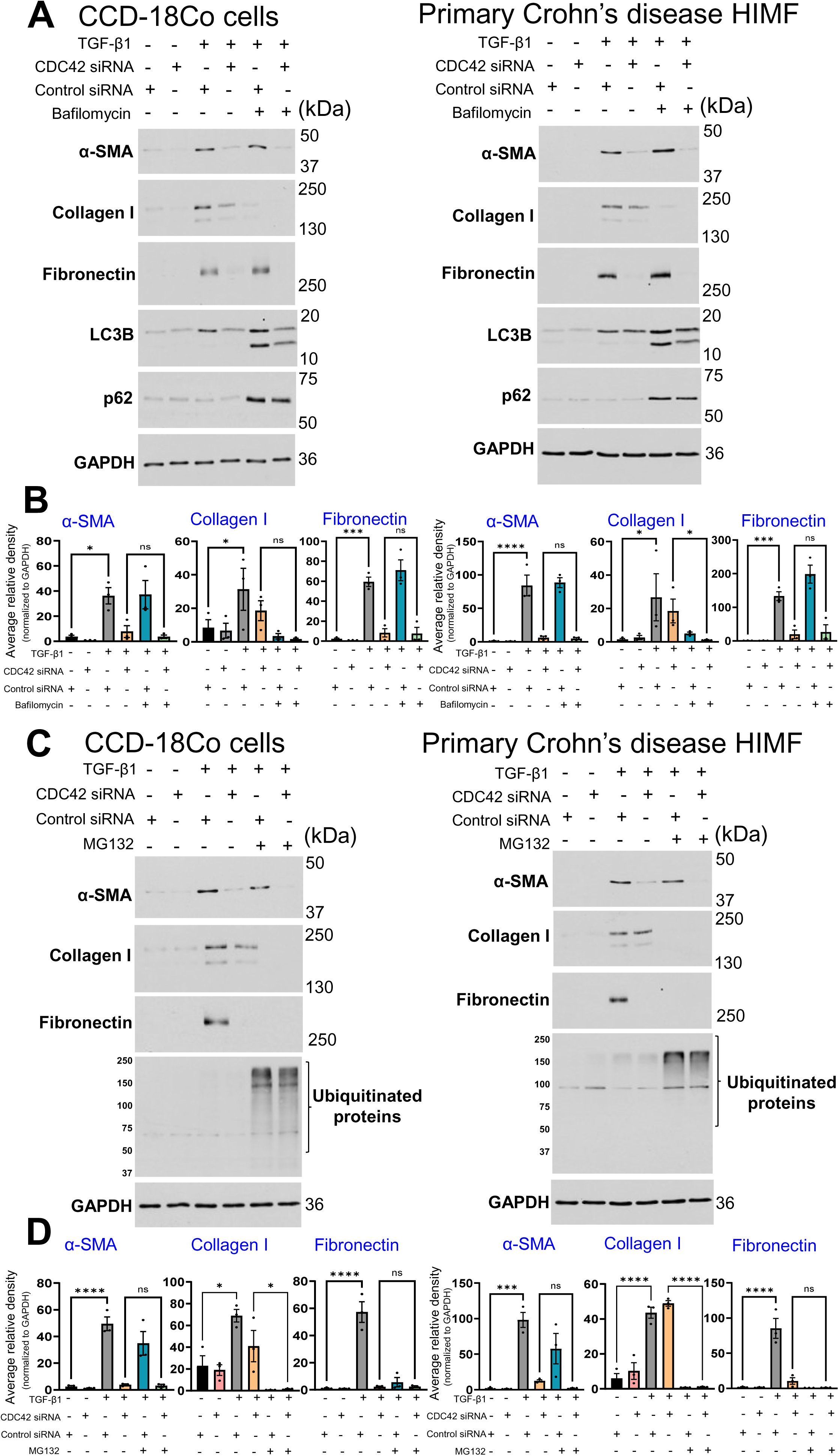
Inhibition of lysosomal and proteasome protein degradation pathways does not restore the expression of cytoskeletal proteins and ECM components in Cdc42-deficient intestinal myofibroblasts. Human CCD-18Co cells and CD HIMF were transfected with control or Cdc42-specific siRNA and stimulated for 48 hours with TGF-β1 in the presence of either a lysosomal inhibitor, bafilomycin A1 (25 nM; **A** and **B**), a proteasomal inhibitor, MG132 (0.5 µM; **C** and **D**) or vehicle. Immunoblotting analysis shows the expression of α-SMA, collagen I and fibronectin. The expression of autophagosomal markers p62 and LC3B (**A**) and poly-ubiquitinated proteins (**C**) were analyzed to demonstrate the efficiency of lysosomal and proteasomal inhibition, respectively. α-SMA, collagen 1 and fibronectin levels **(B, D)** were quantified by densitometry analyses. Average relative band densities represent the values normalized to corresponding GAPDH relative band densities (loading control). Data are shown as mean ± SEM (n=3). *p < 0.05; **p < 0.01; ***p < 0.001 by one-way ANOVA with Tukey’s multiple-comparison test; and *p < 0.05 by one-way ANOVA with Holm-Šídák multiple-comparison test for collagen I expression in CCD-18Co cells, and by the Kruskal-Wallis test followed by Dunn’s post-hoc test for collagen I expression in CD HIMF.

### Cdc42 drives de novo protein translation and mTOR activation

Since our data indicate that loss of Cdc42 attenuates TGF-β dependent activation of intestinal myofibroblasts by mechanisms independent of transcription inhibition and accelerated protein degradation, we next examined a possible role of protein translation. A puromycin incorporation assay was used to examine alterations in global protein translation during fibroblast activation^48,49^. Cells were pulsed with a low dose of puromycin, and the incorporation of puromycin into newly synthesized polypeptides was examined by immunoblotting using anti-puromycin antibody. TGF-β1 exposure of CCD-18Co cells and CD HIMF for 48 h markedly increased puromycin incorporation into different proteins, thereby indicating the increase in protein translation (Figure 6A and B). Remarkably, Cdc42 depletion resulted in a dramatic decrease in the amount of puromycin containing polypeptides both in control and TGF-β1-treated CCD-18Co cells and primary CD HIMF (Figure 6A and B). These data highlight Cdc42 as a key regulator of protein translation both in stimulated and quiescent human intestinal fibroblasts.

**Figure 6.**
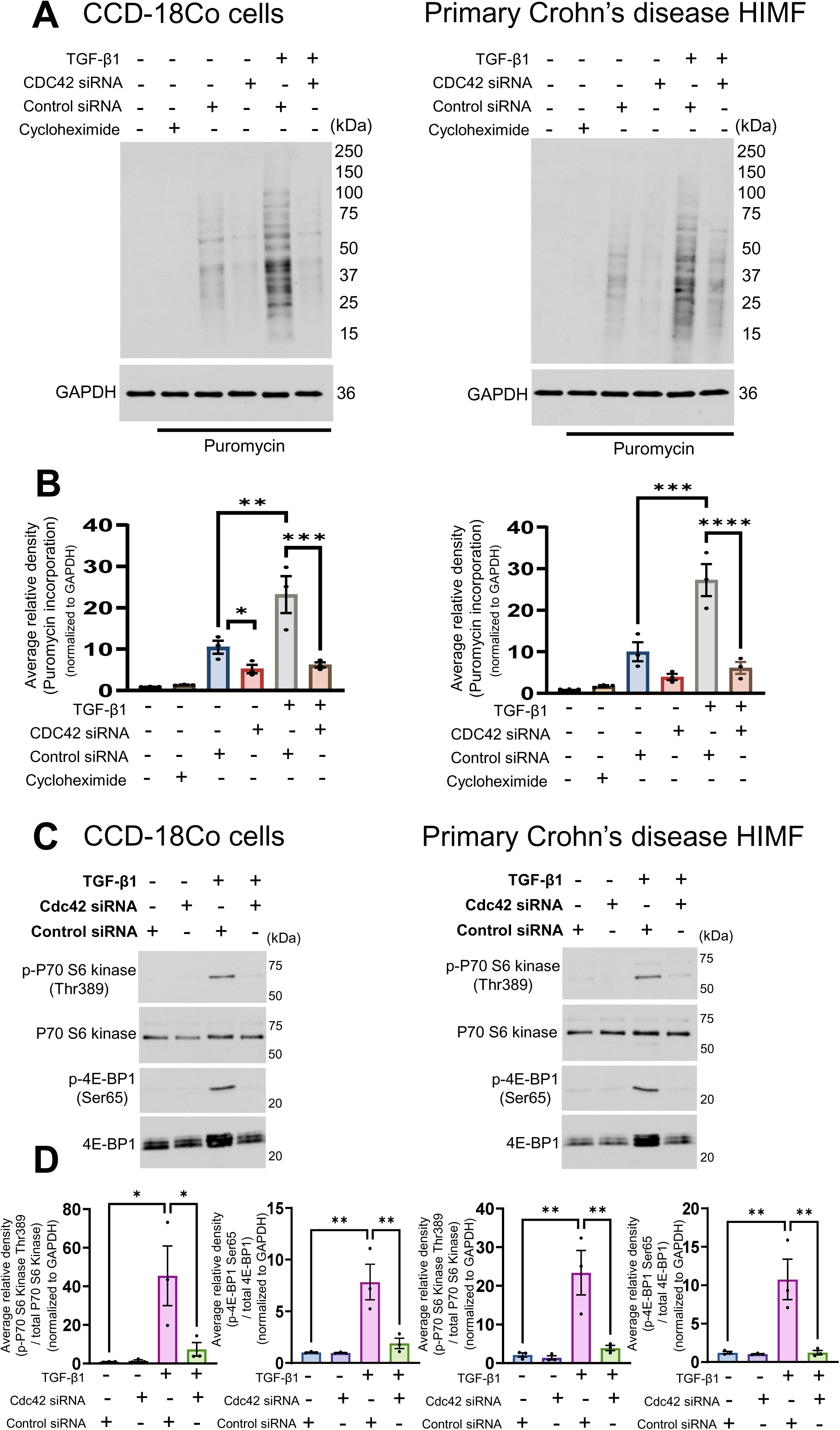
Cdc42 knockdown decreases global protein translation and inhibits mTOR signaling in TGF-β1 activated intestinal myofibroblasts. (**A** and **B**) CCD-18Co cells and CD HIMF were transfected with control or Cdc42-specific siRNAs and stimulated with TGF-β1 for 48 hours. Live cells were pulsed with puromycin for 1 h prior to collection of cell lysates. Immunoblotting analysis with anti-puromycin antibody shows puromycin incorporation in newly synthesized polypeptides. Cell treatment with a classical inhibitor of protein translation, cycloheximide, served as a positive control for the puromycin assay. Puromycin incorporation levels were quantified by densitometry analyses **(B)**. (**C** and **D**) Control or Cdc42-depleted CCD-18Co cells and CD HIMF were stimulated with TGF-β1 for 24 hours. Immunoblotting analysis shows expression of total and phosphorylated forms of major downstream mTOR effectors, p70 S6 kinase and 4E-BP1. Phospho-p70 S6 kinase and phospho-4E-BP1 levels were quantified by densitometry analyses **(D)**. Average relative band densities represent the values normalized to corresponding GAPDH relative band densities (loading control). Data are presented as mean ± SEM (n=3). *p < 0.05 between controls siRNA and cdc42 siRNA incorporated with puromycin groups in CCD-18Co cells by unpaired t-test, and *p < 0.05; **p < 0.01; ***p < 0.001 by one-way ANOVA with Tukey’s multiple-comparison test.

To understand the mechanism by which Cdc42 regulates protein translation, we focused on the key regulator of protein translation, mTOR. mTOR is a known mediator of tissue fibrosis, including intestinal fibrosis^50–57^. Furthermore, a handful of previous studies suggested that Cdc42 could regulate mTOR activity in cancer and epithelial cells^58–61^; however, none of these studies examined the signaling coupling between Cdc42 and mTOR in the context of tissue fibrosis. Expectedly, TGF-β1 stimulation of CCD-18Co cells and primary CD HIMF resulted in mTOR activation, which was manifested by the increased phosphorylation of two major downstream mTOR effectors, p70 S6 kinase and 4E-BP1 (Figure 6C and D). Depletion of Cdc42 significantly decreased TGF-β1 induced phosphorylation of P70 S6 kinase and 4E-BP1, which indicates inhibition of mTOR signaling (Figure 6C and D). Consistently, inhibition of mTOR signaling with rapamycin markedly attenuated the increased expression of α-SMA and major ECM proteins (collagen I and fibronectin) in activated CCD-18Co cells and primary CD HIMF (Suppl. Figure 6). Together, our data demonstrates that Cdc42 stimulates mTOR signaling to upregulate translation of contractile markers and ECM proteins in TGF-β activated human intestinal myofibroblasts.

### Cdc42 regulates fibroblast activation by mechanisms involving AKT, but independently of IQGAP1 and DOCK7

Next, we explored possible mechanisms that link Cdc42 and mTOR activation in TGF-β stimulated human intestinal myofibroblasts. While Cdc42 is known to regulate several signaling pathways that may intercept with mTOR signaling^58–64^, we focused on major downstream Cdc42 effectors previously linked to tissue fibrosis. One is IQGAP, a large scaffolding molecule that is activated by Cdc42^65–70^ and was shown to regulate TGF-β driven fibrogenic epithelial-mesenchymal transition (EMT)^71^ and modulate mTOR activity in cancer cells^62,72^. However, we did not observe increased expression of IQGAP1 protein in TGF-β1 stimulated CCD-18Co cells and primary HIMF (Suppl. Figure 7A). Furthermore, siRNA mediated knockdown of IQGAP1 did not recapitulate the inhibitory effects of Cdc42 depletion on α-SMA and ECM protein expression in activated myofibroblasts (Suppl. Figure 7A).

We also investigated another known Cdc42-associated protein, DOCK7, which plays a dual role as an upstream Cdc42 regulator^73,74^ and a downstream scaffolding protein mediating assembly of a large multiprotein complex encompassing Cdc42 and mTOR^61^. DOCK7 expression appears to be slightly upregulated in TGF-β treated cells (Suppl. Figure 7B). However, siRNA-mediated depletion of DOCK7 did not consistently downregulate expression of α-SMA, collagen I and fibronectin in activated CCD-18Co cells and primary CD HIMF, indicating that DOCK7 is not essential for the fibrogenic activation of human intestinal myofibroblasts (Suppl. Figure 7B). Finally, we examined the role of AKT signaling. While AKT may have multiple roles in regulating TGF-β signaling, a recent report highlights AKT as a component of the multiprotein complex that mediates Cdc42-dependent mTOR activation in breast cancer cells^61^. Consistently, our data demonstrates that AKT is activated by TGF-β in human intestinal myofibroblast cells via Cdc42-dependent mechanisms (Figure 7A,B). Furthermore, a selective AKT inhibitor, MK-2206^75^, efficiently blocked AKT activation and diminished expression of α-SMA and ECM proteins in stimulated CCD-18Co cells and primary CD HIMF (Figure 7C-E). Importantly, MK-2206 also inhibited TGF-β1 induced mTOR activation (Figure 7C,D). Together these data emphasize AKT as an important signaling step during fibrogenic activation of intestinal myofibroblasts and indicate that AKT could serve as signaling transduced from Cdc42 to mTOR to stimulate expression of contractile myofibroblast markers and ECM proteins.

**Figure 7.**
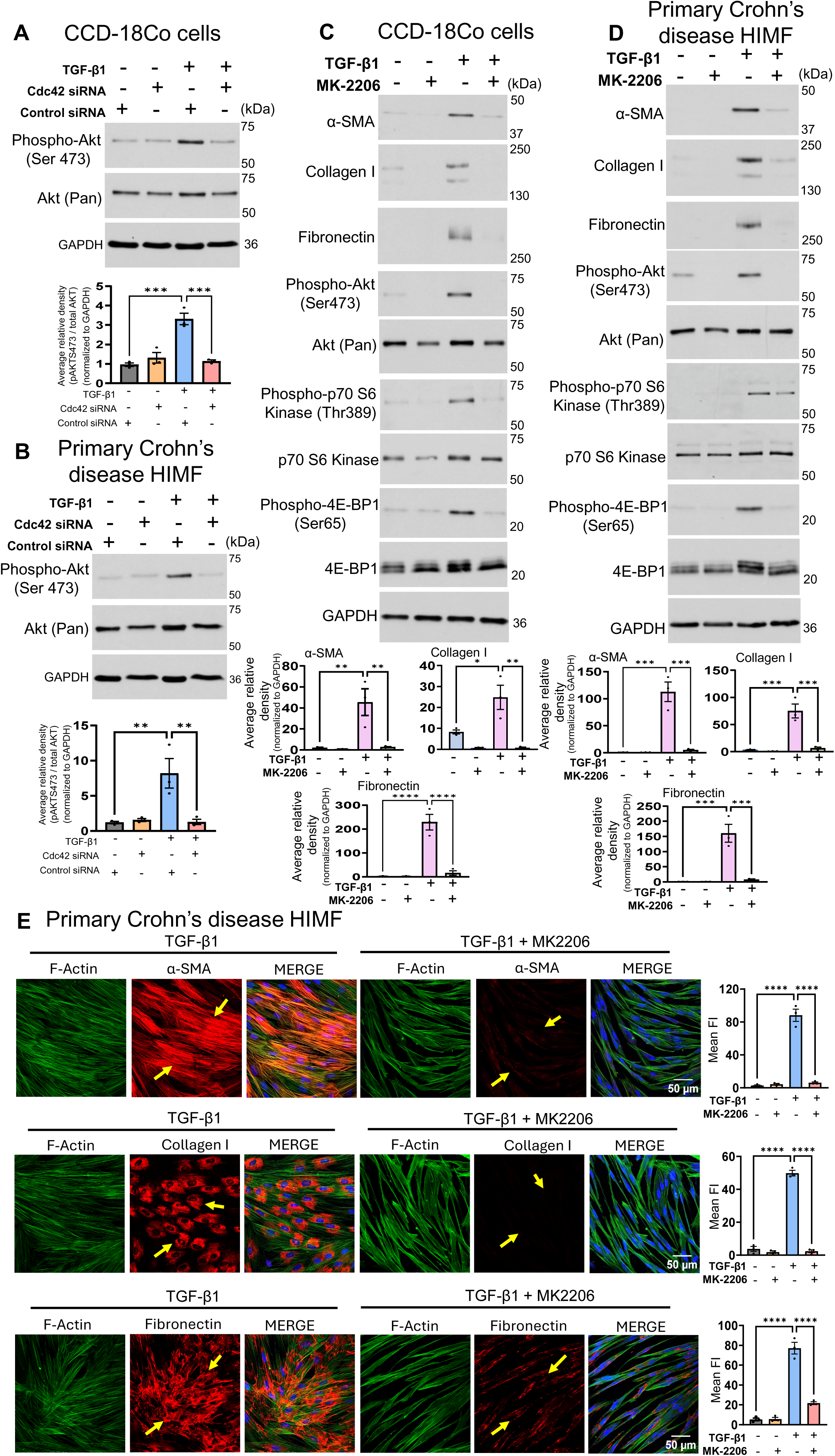
Cdc42 depletion attenuates TGF-β-dependent activation of intestinal myofibroblasts by inhibiting Akt activity. (**A and B**) CCD-18Co cells and HIMF from CD patients were transfected with control or Cdc42-specific siRNA and stimulated with TGF-β1 for 24 hours. Immunoblotting analysis of the expression of total and phosphorylated (active) AKT in total cell lysates. The phospho-AKT levels were quantified by densitometry analyses. (**C-E**) CCD-18Co cells and HIMF from CD patients were stimulated with TGF-β1 for 48 hours in the presence or absence of the AKT inhibitor, MK-2206 (10 µM). (**C and D)** Immunoblotting analysis shows the effects of AKT inhibition on the expression of α-SMA, collagen I, fibronectin, expression of total and phosphorylated Akt, and the expression and activation of downstream mTOR effectors (p70 S6 kinase and 4E-BP1). α-SMA, collagen I and fibronectin levels were quantified by densitometry analyses. Average relative band densities represent the values normalized to corresponding GAPDH relative band densities (loading control). (**E**) Confocal images show the effects of AKT inhibition on α-SMA, collagen I and fibronectin in primary CD HIMF after TGF-β1 stimulation. Images shown are representative of three independent experiments with multiple images taken per slide, and mean fluorescence intensity (FI) of images was measured using the Image J software. Arrows indicate αSMA, collagen I and fibronectin induction in TGF-β1 treated primary CD HIMF and inhibition of αSMA, collagen I and fibronectin in cells stimulated with TGF-β1 in the presence of AKT inhibitor (MK-2206). Data are presented as mean ± SEM (n=3). *p < 0.05; **p < 0.01; ***p < 0.001 by one-way ANOVA with Tukey’s multiple-comparison test. Scale bars, 50 µm.

## Discussion

In this study, we identified a novel signaling mechanism that regulates activation of intestinal myofibroblasts by a key profibrotic factor TGF-β. This mechanism involves a major small GTPase, Cdc42, that activates a downstream AKT-mTOR signaling module to drive the pro-fibrotic transformation of the fibroblast phenotype.

The role of Cdc42 in profibrotic HIMF activation is supported by our findings that either pharmacologic inhibition or genetic depletion of this small GTPase markedly attenuated TGF-β-induced expression of contractile cytoskeletal and ECM proteins. Consistently, Cdc42 inhibition blocked ECM secretion by the activated myofibroblasts. This data strongly suggests that Cdc42 controls two major aspects of phenotypic plasticity in activated HIMF, such as increased ECM production/secretion and enhancement of the contractile actin cytoskeleton. The discovered mechanism is novel since Cdc42 has not been previously implicated in HIMF activation or intestinal fibrosis. This mechanism may not be limited to the intestine but also mediate stromal cell activations in other organs, such as kidneys and liver where chronic inflammation and fibrosis are associated with Cdc42 activation^32,33,35,36^.

While dissecting the place for Cdc42 within the TGF-β stimulated signaling cascade we found this GTPase does not regulate signaling from TGF-β receptor and transcription of TGF-β dependent genes. These findings are consistent with a previous study of differentiated mesenchymal stem cells, where loss of Cdc42 affected neither TGF-β induced SMAD activation, nor transcription of α-SMA and collagen I^24^. We also did not find evidence that Cdc42 controls degradation of ECM and contractile cytoskeletal proteins. Instead, our data suggests that Cdc42 regulates protein translation in intestinal myofibroblasts. Indeed, according to the puromycin incorporation assay, loss of Cdc42 markedly inhibited de novo protein synthesis under resting conditions and in TGF-β activated cells. The observed inhibitory effects are very robust and involve multiple polypeptides, which signifies Cdc42 as a key regulator of global protein translation. This Cdc42 function remains surprisingly overlooked with a very limited number of publications that described regulation of protein translation by known downstream molecular effectors of Cdc42 in cancer cells^76,77^. Importantly, protein translation is an emerging mechanism that drives ECM secretion in activated myofibroblasts. In this regard, we recently found that activation of the translation initiation machinery and increased polysomal accumulation of fibronectin mRNA underlie the increased secretion in HIMF activation by bacterial flagellin^78^.

Another important finding of our study suggests that a master-regulator of protein translation, mTOR, acts downstream of Cdc42 in activated HIMF to drive TGF-β induced expression of contractile cytoskeletal and ECM proteins. mTOR is a serin/threonine kinase which plays major role in controlling cell growth and metabolism^79–81^. mTOR regulates eukaryotic mRNA translation initiation by two parallel mechanisms: phosphorylation and activation of the p70 ribosomal S6 kinase (p70-S6K) and phosphorylation and inactivation of eukaryotic translation initiation factor 4E-binding protein (4E-BP)^80,81^. While key roles of mTOR in regulating tissue fibrosis have been previously investigated^50–57^, upstream signaling events that promote mTOR function in activated myofibroblast remain incompletely understood. Our study provides the first evidence that Cdc42 stimulates mTOR to drive profibrotic activation of human myofibroblasts. Such activation is reflected by Cdc42-dependent modulation of both mTOR effector phosphorylation, p70-S6K and 4E-BP, in TGF-β-stimulated HIMF. Consistently, pharmacologic inhibition of mTOR with rapamycin potently blocked profibrotic activation of human intestinal myofibroblasts. While our findings are limited to myofibroblast activation *in vitro* it is tempting to suggest that the Cdc42-mTOR signaling module could be also important for tissue fibrosis *in vivo*.

What are the mechanisms linking Cdc42 and mTOR signaling in activated myofibroblasts? While the signaling interplay between mTOR and Cdc42 has been previously described in epithelial and cancer cells^58–61^ the underlying molecular mechanisms remain poorly understood. Since Cdc42 was shown to poorly interact with mTOR complexes^82^, the Cdc42-dependent stimulation of mTOR could be mediated either by specific scaffolding proteins or intermediate signaling events^59,61^. However, our data argues against the roles of two important Cdc42 scaffolding proteins, IQGAP1 and DOCK7 in HIMF activation. By contrast, our data supports the intermediate signaling connections between Cdc42 and mTOR and specifically highlights the possible roles of AKT signaling. AKT is a well-known mTOR regulator that targets the tuberous sclerosis complex 2 (TSC2) protein serving as an upstream inhibitor of mTOR^83–85^. AKT-dependent phosphorylation of TSC2 relieves mTOR inhibition. We observed that AKT is activated in TGF-β stimulated HIMF and such activation depends on Cdc42. Furthermore, the inhibitory analysis demonstrated that AKT positively regulates mTOR activity and is essential for production of contractile markers and ECM proteins in activated HIMF. While we did not investigate the mechanisms by which Cdc42 could stimulate AKT, at least two major downstream effectors of Cdc42 are known to regulate AKT activity in different cancer cells^86–88^. One such effector is a p21 kinase (PAK) protein family that is known to positively regulate AKT activity via phosphorylation or scaffolding mechanisms^89–91^. The second effector is an activated Cdc42-associated kinase 1 (Ack1), a non-receptor tyrosine kinase which is activated by various growth factor receptor tyrosine kinases and directly phosphorylates AKT to drive cancer cell proliferation and motility^92–94^. Future studies are needed to establish which Cdc42 effector kinases are responsible for mTOR stimulation and phenotypic plasticity of activated HIMF.

Since Cdc42 has been implicated in tumorigenesis and other chronic disorders, several pharmacological inhibitors have been developed to target this small GTPase in human diseases^95–97^. Our demonstration of the critical role of Cdc42 in profibrotic activation of HIMF raises an important question whether Cdc42 inhibitors could be used as novel therapeutic agents to treat intestinal fibrosis in IBD patients. While this question awaits to be addressed by direct experiments, a prolonged global Cdc42 inhibition may have detrimental effects due to important functions of this small GTPase in intestinal epithelium. Indeed, a selective deletion of Cdc42 in murine intestinal epithelium was shown to damage intestinal epithelial architecture resulting in crypt dysplasia, villous shortening and spontaneous inflammation^98^. Likewise, a selective loss of Cdc42 in the intestinal stem cell compartment caused dramatic hyperproliferation and loss of epithelial apico-basal cell polarity^99^. Such detrimental effects of cellular Cdc42 deletion suggest that a more feasible approach would be to target downstream effectors of Cdc42 that are essential for its profibrotic signaling, in order to develop novel anti-fibrotic therapies.

In summary, our study identifies a novel signaling mechanism that drives profibrotic activation of human intestinal myofibroblasts. The mechanism involves stimulation of Cdc42 small GTPase that signals to the AKT-mTOR molecular module to activate translation of ECM and contractile cytoskeletal proteins. Pharmacological targeting of this signaling cascade may have potential for developing novel antifibrotic strategies.

### Limitations of the study

While our genetic and pharmacological approaches in this study provided proof-of-principle that Cdc42 functions as a key regulator of intestinal fibrosis via activation of the AKT-mTOR signaling module; several limitations should be acknowledged - (i) studies using animal models are needed to recapitulate the complex-cellular interactions, inflammatory milieu and extracellular matrix remodeling that occur during intestinal fibrogenesis in vivo to confirm the role of CDC42 and evaluate therapeutic potential of its inhibition in intestinal fibrosis; (ii) limited number of patient-derived samples may not capture the heterogeneity of intestinal fibrosis across patient populations; and (iii) the plausible mechanisms linking Cdc42 and Akt-mTOR stimulation in activated myofibroblasts remains an open question that warrants further investigations, (iv) and lastly the mechanisms of Cdc42 activation in intestinal myofibroblasts by profibrotic stimuli have not been studied and require further research.

## RESOURCE AVAILABILITY

### Lead contact

Further information and requests for resources and reagents should be directed to and will be fulfilled by the lead contact, Andrei Ivanov.

### Materials availability

Reagents generated in this study will be made available upon request but may require a completed materials transfer agreement.

### Data and code availability

- No code was generated in this study.
- Any additional information required to reanalyze the data reported in this paper is available from the lead contact upon request.

## Acknowledgments

The authors thank the Ivanov’s and Rieder’s lab for helpful discussions. Confocal microscopy performed at the Cleveland Clinic Research Digital Imaging Microscopy Core utilized the Leica SP8 confocal microscope that was purchased with funding from the National Institutes of Health SIG grant 1S10OD019972-01. This work was supported by National Institutes of Health grant (NIH)-NIDDK grant R01 DK132038 and a Synergy Award from the Kenneth Rainin Foundation to F.R. and A.I.I. We also deeply appreciate the contribution of CD patients who donated the tissues for isolation of primary HIMF.

## Author contributions

Conceptualization, A.I. and A.Z.; methodology, A.I., A.Z. and G.C; scRNAseq analysis, P.K.M.; confocal microscopy: A.Z., A.M.M and N.G.N.; isolation of primary HIMF from CD patients: G.C. and Y.W.; validation, A.I., A.Z., G.C. and P.K.M; formal data analysis, A.Z., G.C., P.K.M. and N.G.N; investigation, A.I.I., A.Z., G.C. and P.K.M; resources and intellectual input, A.I.I and F.R.; data curation, A.I.I and A.Z.; writing—original draft preparation, A.I.I.; writing—review and editing, A.I.I., F.R., A.Z. and P.K.M; visualization and supervision, A.I.I.; project administration, A.I.I.; funding acquisition: A.I.I. and F.R. All authors have read and agreed to the published version of the manuscript.

## STAR★METHODS

### Key resource table

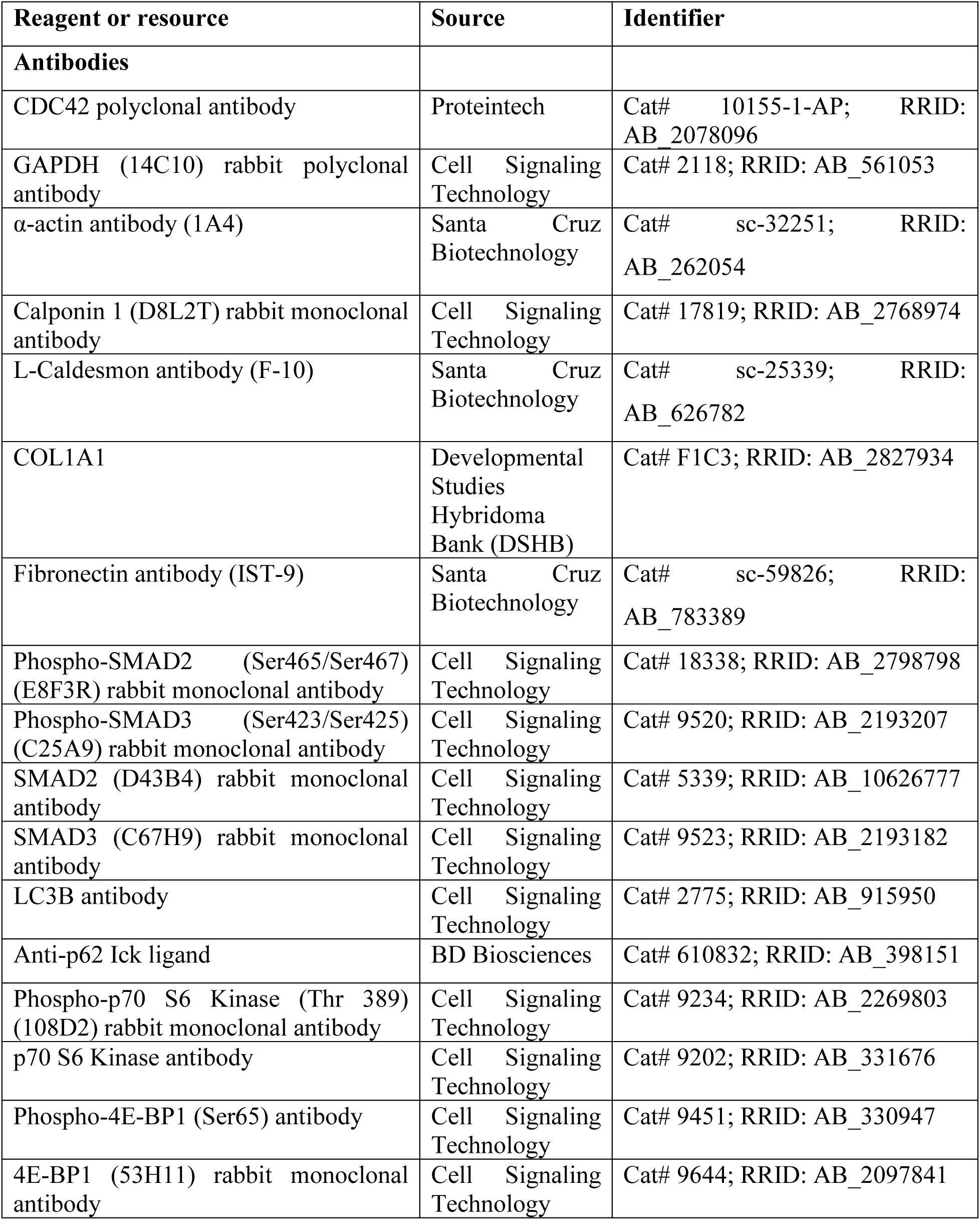
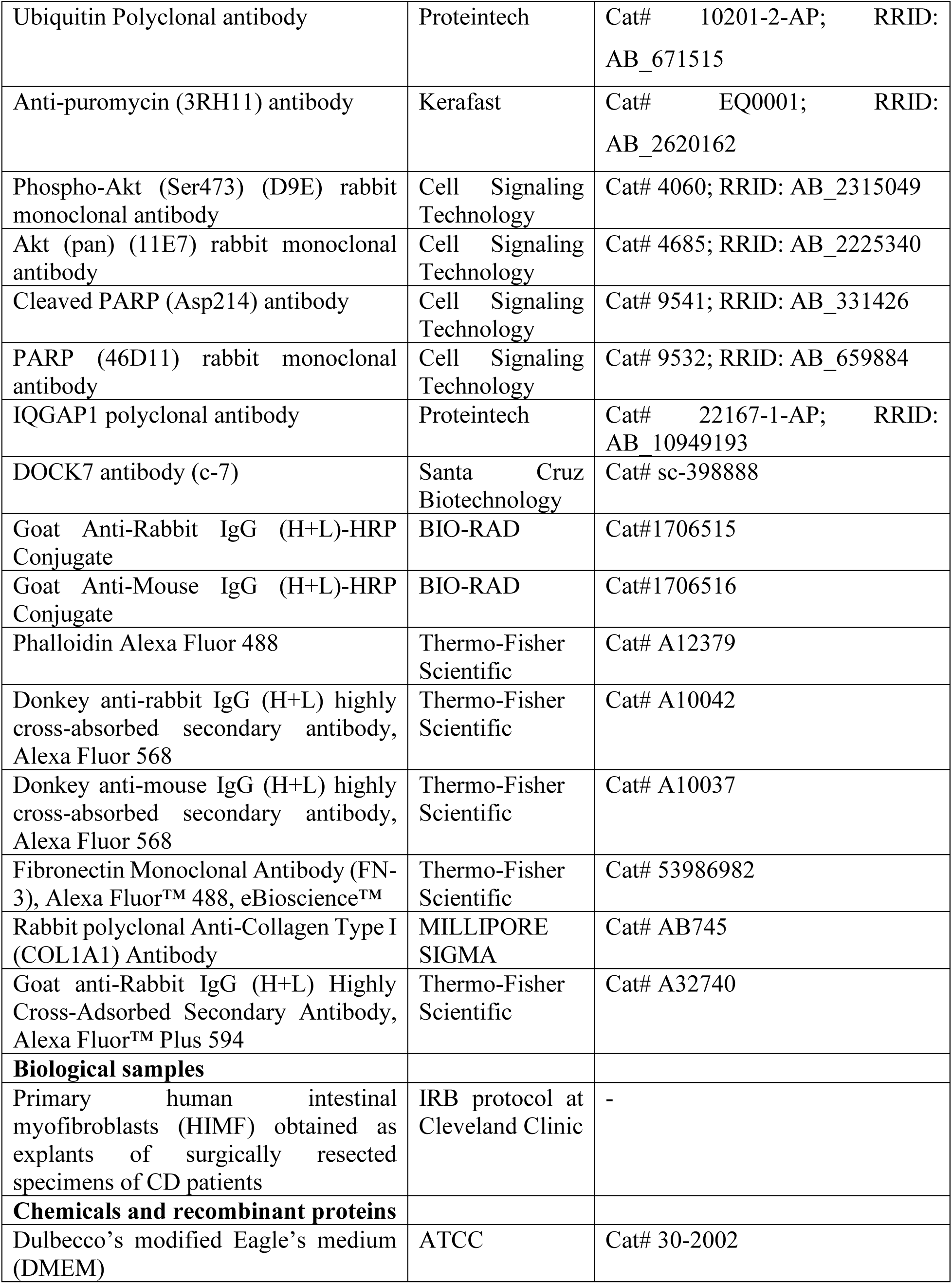
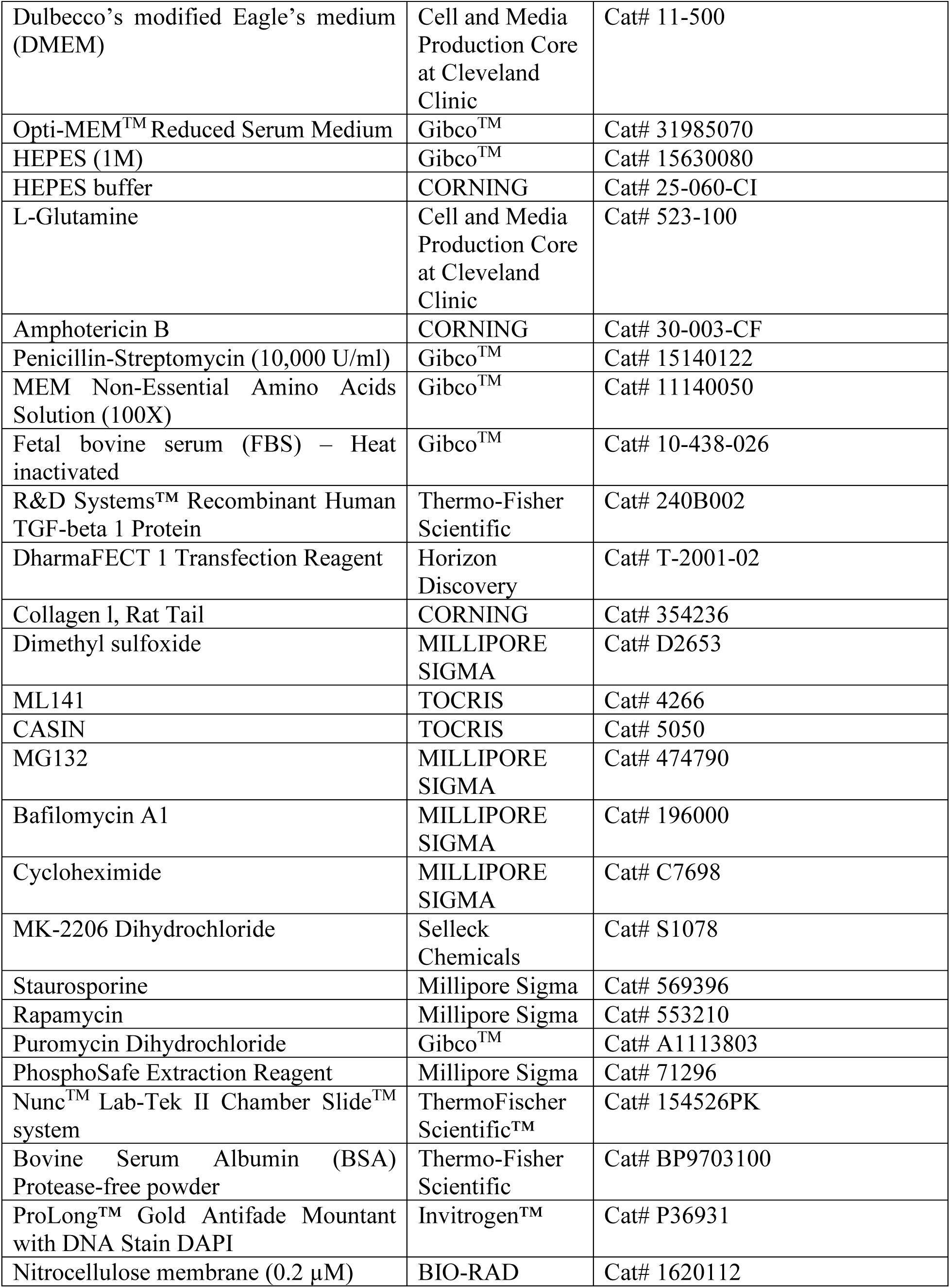
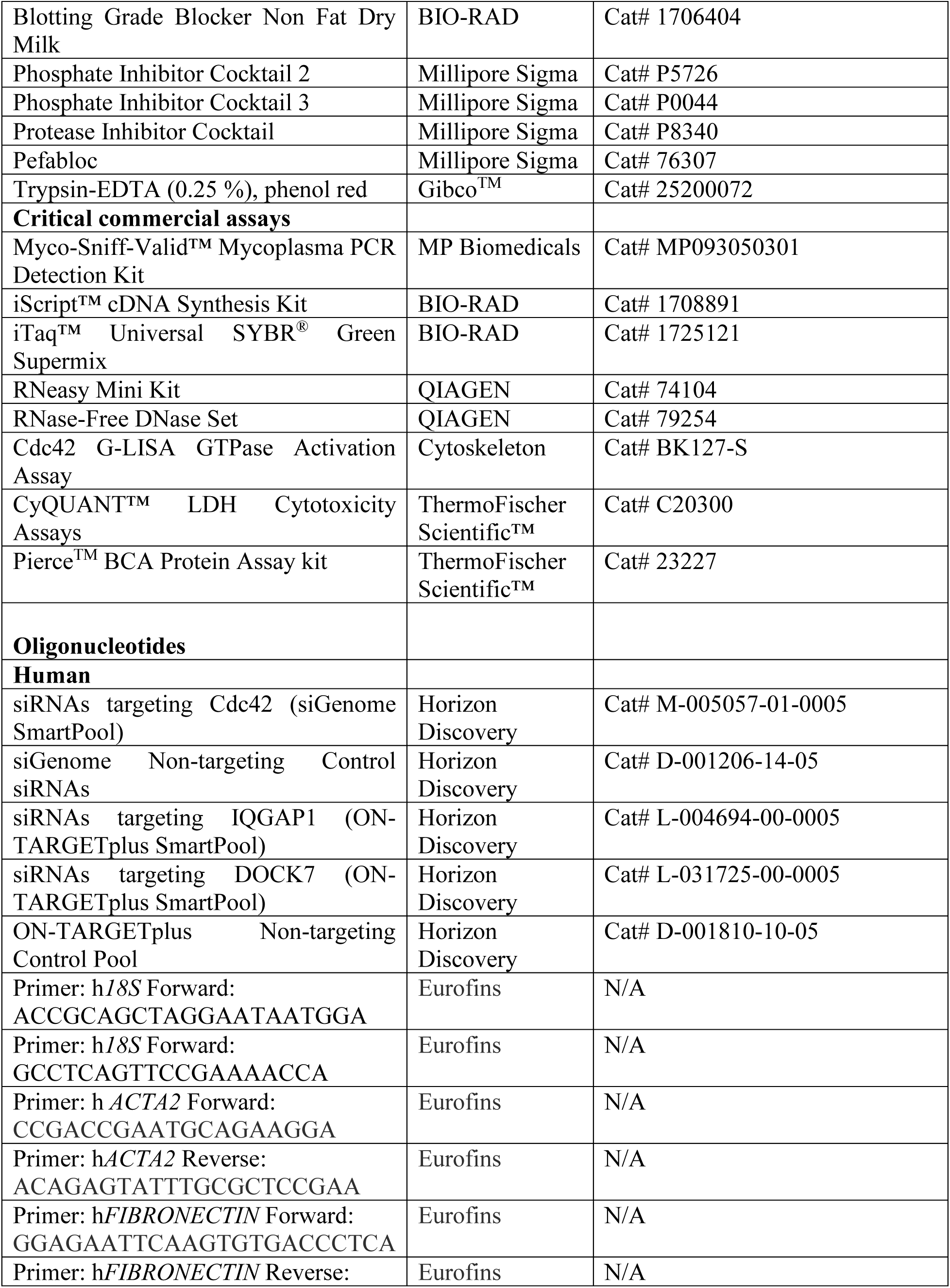
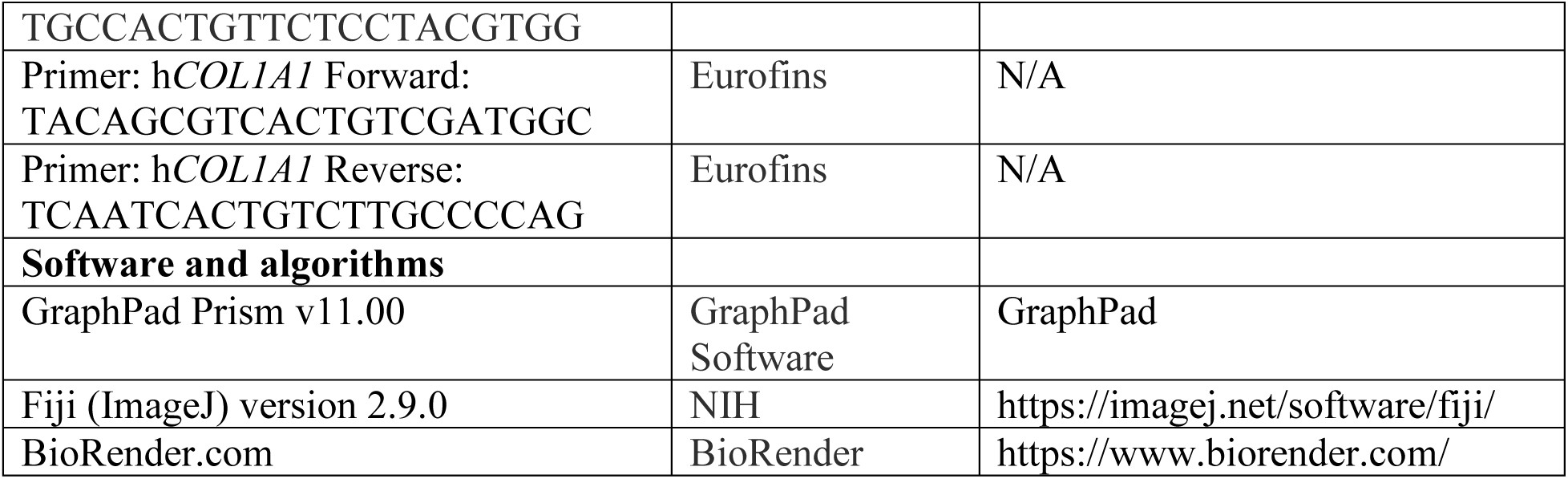

### Experimental model and study participant details

#### Cell lines

Human cell line normal colonic fibroblast CCD-18Co (CRL-1459) were obtained from the American Type Culture Collection (ATCC, Manassas, VA, USA) and cultured in Dulbecco’s modified Eagle’s medium (DMEM), high glucose (ATCC, Cat# 30-2002) supplemented with 10% fetal bovine serum (FBS), 1M HEPES and minimal essential medium (MEM) non-essential amino acids and penicillin-streptomycin (10,000 U/ml) at 37°C and 5% CO_2_. Cell line was used for no more than 6 passages and were confirmed to be mycoplasma negative by PCR testing.

#### Primary human intestinal myofibroblasts

Surgical resection specimens were used for isolation of primary human intestinal myofibroblasts (HIMF) as previously described^37,78^. This protocol was approved by the Cleveland Clinic Institutional Review Board (number 19-1035). HIMF were obtained as explants of surgically resected intestinal mucosa of CD patients, grown to subconfluence in DMEM (Cell and Media Production Core at Cleveland Clinic) supplemented with 10% fetal bovine serum (FBS), HEPES (Corning), L-glutamine, penicillin-streptomycin and amphotericin B, and established as long-term cultures fed twice weekly and subcultured at confluence^37,78^. For all the experiments involving primary cells, HIMF were derived from the colon or ileum of CDi or CDs patients depending on tissue availability and these HIMF were used interchangeably under one category as ‘CD’. Primary HIMF isolated from both male and female donors were used in the study.

### Method

#### details Reagents

Chemicals and the working concentrations used in the pharmacological assays are as follows: ML141 (Tocris, 4266, 5 or 10 µM), Casin (Tocris, 5050, 5 µM), MG132 (Millipore Sigma, 474790, 0.5 µM), Bafilomycin A1 (Millipore Sigma, 196000, 25 nM), cycloheximide (Millipore Sigma, C7698, 20 µM), MK-2206 dihydrochloride (Selleck Chemicals, S1078, 10 µM), staurosporine (Millipore Sigma, 569396, 100 nM), rapamycin (Millipore Sigma, 553210, 2.5 or 5 µM), dimethyl sulfoxide (DMSO) (Millipore Sigma, D2653). Antibodies were purchased from Cell Signaling Technologies (Danvers, MA, USA) including calponin 1 (17819), phosphor-SMAD2 (Ser465/Ser467) (18338), phosphor-SMAD3 (Ser423/Ser425) (9520), SMAD2 (5339), SMAD3 (9523), LC3B (2775), phosphor-P70 S6 kinase (Thre389) (9234), p70 S6 Kinase (9202), phosphor-4E-BP1 (Ser65) (9451), 4E-BP1 (9644), phosphor-Akt (Ser473) (4060), Akt (pan) (4685), cleaved PARP (Asp214) (9541) and PARP (9532), GAPDH (2118), Santa Cruz Biotechnologies (Santa Cruz, CA, USA) including α-actin (sc-32251), L-caldesmon (sc-25339), fibronectin (sc-59826), DOCK7 (sc-398888), Proteintech including Cdc42 (10155-1-AP), ubiquitin (10201-2-AP), IQGAP1 (22167-1-AP), collagen 1, p62 and puromycin antibodies were procured from developmental studies hybridoma bank-DSHB, BD Biosciences and Kerafast, respectively, Bio-Rad (Hercules, California) including HRP-conjugated anti-rabbit IgG, 1706515 and HRP-conjugated anti-mouse IgG, 1706516, and ThermoFisher Scientific including Phalloidin Alexa Fluor 488, Donkey anti-rabbit IgG (H+L) highly cross-absorbed secondary antibody, Alexa Fluor 568, Donkey anti-mouse IgG (H+L) highly cross-absorbed secondary antibody, Alexa Fluor 568, Fibronectin Monoclonal Antibody (FN-3), Alexa Fluor™ 488, eBioscience™ and Goat anti-Rabbit IgG (H+L) Highly Cross-Adsorbed Secondary Antibody, Alexa Fluor™ Plus 594. siRNAs targeting Cdc42 (M-005057-01-0005), IQGAP1 (L-004694-00-0005), DOCK7 (L-031725-00-0005), and non-targeting control (D-001206-14-05 and D-001810-10-05) siRNAs were obtained from Horizon Discovery (Cambridge, UK).

#### Single cell RNA sequencing (scRNA-seq)

Single cell RNAseq data were reanalyzed from data generated in our prior publication^37^. Data were analyzed using R version 4.5.3 and the following R packages to create the dot plots and bar charts shown in Figure 1 and supplementary figures 1 and 2. Following R packages were used: AnnotationDbi v. 1.72.0, Biobase v. 2.70.0, BiocGenerics v. 0.56.0, clusterProfiler v. 4.18.4, expss v. 0.11.7, generics v. 0.1.4, ggside v. 0.4.1, IRanges v. 2.44.0, kableExtra v. 1.4.0, maditr v. 0.8.7, Nebulosa v. 1.20.1, openxlsx v. 4.2.8.1, org.Hs.eg.db v. 3.22.0, paletteer v. 1.7.0, patchwork v. 1.3.2, rwantshue v. 0.0.3, S4Vectors v. 0.48.0, scCustomize v. 3.2.4, SCpubr v. 3.0.1, Seurat v. 5.5.0, SeuratObject v. 5.2.0, sp v. 2.2.1, tictoc v. 1.2.1, tidytext v. 0.4.3, tidyverse v. 2.0.0.

#### Cdc42 G-ELISA activation assay

Activation of Cdc42 was performed using a G-LISA® based Cdc42 activation assay kit (Cytoskeleton, Inc, BK127) per manufacturer’s instructions. Briefly, normal human colonic fibroblast CCD-18Co cells and primary human intestinal myofibroblasts (HIMF) isolated from CD patients were seeded in 6-well plates at a density of 2 × 10^5^ cells per well and allowed to grow for 2 days. Subsequently, cells were serum-deprived overnight and then stimulated with TGF-β1 (5 ng/ml) for time points indicated in the corresponding figure. After TGF-β1 stimulation, cells were washed in 1X ice-cold DPBS (Gibco, 14040133), lysed in ice-cold lysis buffer (part #GL535) containing protease inhibitor cocktail and cleared at 10,000 x g for 1 min at 4°C. Protein concentrations of individual samples were determined using Pierce BCA protein assay kit (Thermo Scientific, 23227). Protein concentrations were normalized across samples by addition of required amount of ice-cold lysis buffer to equalize all concentrations. Normalized protein samples along with lysis buffer (blank) and CDCA (constitutively active Cdc42 control protein) were then added to equilibrated microplate containing Cdc42-GTP strips and incubated at 4°C for 15 min on orbital microplate shaker (400 rpm). Subsequently plates were washed with wash buffer and then incubated with antigen presenting buffer for 2 min at room temperature. Then, diluted Cdc42 primary antibody was added, and the plate was incubated for 30 min in the orbital microplate shaker (400 rpm) at room temperature. Following this, diluted secondary antibody was added and leave the plate on a microplate shaker (400 rpm) at room temperature for 30 min. Finally, the mixed HRP A/B was added, incubated at room temperature for 15 min, and the reaction was then stopped by adding HRP stop buffer. The absorbance values were measured using a microplate spectrophotometer at 490 nm.

#### Treatment of intestinal fibroblasts with pharmacological inhibitors

For experiments, CCD-18Co cells and primary HIMF were seeded in 6-well plates and allowed to grow for 2-3 days. Cells were serum deprived overnight and subsequently stimulated with 5 ng/ml recombinant TGF-β1 (240-B-002, R&D Systems) with or without Cdc42 inhibitors, ML141 or Casin at indicated concentrations for 48 hours. In addition, non-stimulated cells were treated with these inhibitors for 48 hours. To assess the functional importance of Akt signaling, CCD-18Co cells and primary HIMF were stimulated with TGF-β1 for 48 hours in the presence or absence of MK-2206 dihydrochloride (Akt inhibitor) (Selleck Chemicals, 10 µM). In all described above experiments, the vehicle control cells were treated with 0.1% DMSO.

#### Small-interference RNA-mediated transfections

Small interference siRNA-mediated transient knockdown of Cdc42 or IQGAP1 or DOCK7 in CCD-18Co cells and primary HIMF was carried out using specific small interfering RNA (siRNA) and scrambled control siRNA (Horizon Discovery, Cambridge, United Kingdom). Briefly, CCD-18Co cells and primary HIMF were seeded in 6-well plates and allowed to adhere overnight. Cells were transfected using DharmaFECT1 transfection reagent (Horizon, Cambridge, T-2001–02) in Opti-MEM I medium (Thermo Fisher Scientific, 31985070) according to the manufacturer’s protocol, with final siRNA concentrations of 25 or 50 nM. After transfection for 8 hours, the Opti-MEM I medium was replaced with regular DMEM culture medium for 48 h. Next, cells were serum-deprived overnight and stimulated with TGF-β1 for the indicated times and subjected to either total protein lysate preparation or RNA extraction for subsequent analysis.

#### Extracellular matrix deposition assay

Deposition of extracellular matrix (ECM) by intestinal myofibroblasts was assayed using a method as previously described^37,100^. Briefly, primary HIMF were plated in a 96-well dark-walled imaging plate (Greiner, BioOne, 655090) and allowed to attach overnight. Cells were then treated with Cdc42 inhibitors (ML141 or Casin) at indicated concentrations with or without TGF-β1 in serum-free medium for 5 days to allow ECM accumulation. Separately, primary HIMF were transfected with Cdc42-specific small interfering RNA (siRNA) and scrambled control siRNA in 6-well plate, later trypsinized and plated in 96 well dark-walled imaging plate. Cells were then treated with or without TGF-β1 in serum-free medium for 5 days. At the end of the treatment, the treated wells were washed with PBS and decellularized with ammonium hydroxide. Deposited ECM was fixed by exposure to absolute methanol at −20°C and stained with Alexa Fluor 488-conjugated anti-fibronectin (EBioscience, clone FN-3, 1:500 dilution) or rabbit polyclonal anti-collagen type I (COL1A1) antibody (Millipore Sigma, Burlington, MA, 1:100 dilution) followed by secondary Alexa Fluor-594-conjugated goat anti-rabbit at 1:1000 dilution (ThermoFischer Scientific™). Fluorescence intensities and images were captured using a Cytation5 scanner (Agilent Biotek).

#### Protein degradation pathways analysis

CCD-18Co cells and primary HIMF were plated in 6-well plates and allowed to adhere overnight. Subsequently, the cells were transfected with either Cdc42-specific siRNA or control siRNA, and were serum-deprived at 48 hours post-transfection. Serum deprived cells were stimulated with TGF-β1 for 24 hours followed by re-addition of TGF-β1 in the presence of either bafilomycin A1 (Baf A_1_) (25 nM), MG132 (0.5 µM), or vehicle. The inhibition of targeted lysosomal and proteosomal degradation pathways was validated by analyses of expression levels of autophagosomal markers p62 and LC3B and poly-ubiquitinated proteins, respectively.

#### Gel electrophoresis and immunoblotting

CCD-18Co cells and primary HIMF were scraped and homogenized in RIPA buffer (20 mM Tris, 150 mM EDTA, 2 mM EDTA, 2 mM EGTA, 1% sodium deoxycholate, 1% Triton X-100 (TX-100, 0.1% SDS, pH 7.4) containing phosphatase inhibitor cocktails 2 and 3, protease inhibitor cocktail and Pefabloc^®^ (Millipore-Sigma, Cat# P5726, P0044, P8340 and 76307, respectively). Cell lysates were then cleared by centrifugation at 14,000 x g for 20 minutes to pellet debris and supernatants were collected, diluted with 4x Laemmli sample loading buffer along with IM dithiothreitol (DTT) and boiled. The protein concentrations in each lysate sample were estimated using the Pierce BCA protein assay kit (ThermoFischer Scientific™) according to manufacturer’s instructions. Equal amounts of cellular proteins (3-5 µg) were separated using SDS-polyacrylamide gels. Following this, the proteins were transferred to nitrocellulose membrane and the membranes were then blocked with 3% bovine serum albumin (BSA) or 5% non-fat dry milk in TBS-T (Tris-buffered saline with Tween 20) for 1 hour at room temperature. These blots were further incubated with primary antibodies diluted in 3% bovine serum albumin (BSA) or 5% non-fat dry milk in TBST-T for 1 hour or overnight incubation. After primary antibody incubation, membranes were washed with TBS-T and then the blots were incubated for 1 hour at room temperature in horseradish peroxidase-conjugated secondary antibodies at 1:5000 dilution. Subsequently, the immunolabelled proteins in blots were visualized using enhanced chemiluminescence Western Blotting Substrate (Cytiva Amersham ECL Western Blotting Detection Reagent, RPN2209; Millipore Sigma – Immobilon Classico and Forte Western HRP substrate, WBLUC0100 and WBLUF0100, respectively). Band intensities were quantified by densitometry using Image J software (National Institute of Health, Bethesda, MD). The primary antibodies were used at the following dilutions: CDC42 (rabbit, 1:1000, Proteintech); α-SMA (mouse, 1:1000, Santa Cruz Biotechnology); CALPONIN 1 (rabbit, 1:1000, Cell Signaling Technology); L-CALDESMON (mouse, 1:1000, Santa Cruz Biotechnology); COL1A1 (mouse, 1:500, DHSB); FIBRONECTIN (mouse, 1:200, Santa Cruz Biotechnology); p-SMAD2 (rabbit, 1:500, Cell Signaling Technology); p-SMAD3 (rabbit, 1:500, Cell Signaling Technology); SMAD2 (rabbit, 1:1000, Cell Signaling Technology); SMAD3 (rabbit, 1:1000, Cell Signaling Technology); UBIQUITIN (rabbit, 1:1000, Proteintech); LC3B (rabbit, 1:500, Cell Signaling Technology); p62 (mouse, 1:1000, BD Biosciences); p-P70 S6 Kinase (rabbit, 1:500, Cell Signaling Technology); P70 S6 Kinase (rabbit, 1:1000, Cell Signaling Technology); p-4E-BP1 (rabbit, 1:500, Cell Signaling Technology); 4E-BP1 (rabbit, 1:1000, Cell Signaling Technology); PUROMYCIN [3RH11] (mouse, 1:500, Kerafast); p-AKT (Ser473) (rabbit, 1:500, Cell Signaling Technology); AKT (PAN) (rabbit, 1:1000, Cell Signaling Technology); Cleaved PARP (rabbit, 1:1000, Cell Signaling Technology); PARP (rabbit, 1:1000, Cell Signaling Technology); IQGAP1 (rabbit, 1:500, Proteintech); DOCK7 (mouse, 1:500, Santa Cruz Biotechnology); GAPDH (rabbit, 1:2000, Cell Signaling Technology).

#### RNA extraction and quantitative RT-PCR

Total RNA was isolated from indicated samples using the RNeasy Mini kit (Qiagen, 74104) according to manufacturer’s instructions. RNA was reverse transcribed using iScript reverse transcriptase in the presence of random hexamers and oligo-dT primers (Bio-Rad, 1708891). Real-time quantitative PCR was performed using iTaq™ Universal SYBR^®^ Green Supermix (Bio-Rad, 1725121) on CFX96 Real-Time system in the presence of gene-specific primers (*18S, ACTA2, FIBRONECTIN, COL1A1*). The primer sequences used to determine gene expressions are listed in key resources table.

#### Cell cytotoxicity measurement

CCD-18Co cells and primary HIMF were seeded in 96-well plates and allowed to grow for 2 days. Following this, the cells were serum-deprived overnight and treated with Cdc42 inhibitors (ML141 or Casin) in the presence or absence of TGF-β1 for 48 hours. In addition, cells were transfected with either Cdc42-specific siRNA or control siRNA, serum deprived and stimulated with or without TGF-β1 for 48 hours. Lactate dehydrogenase activity (LDH) assay was performed using a CyQUANT™ LDH Cytotoxicity Assay Kit (ThermoFischer Scientific) according to manufacturer’s instructions. Briefly, cell-culture medium from each sample of above indicated experiments and culture medium from cells lysed with 10X lysis buffer (from kit, maximum LDH activity) were collected to a new 96-well culture plate and then reaction mixture was added and the plate was incubated for 30 minutes at room temperature. Next, stop solution was added and absorbance was measured at 490 nm and 680 nm using a microplate spectrophotometer. The percentage cytotoxicity was calculated using the following formula: [LDH activity of treated sample – spontaneous LDH activity) / (Maximum LDH activity – Spontaneous LDH activity] × 100. Spontaneous LDH refers to LDH activity of cells treated with sterile water only. Furthermore, cell cytotoxicity following Cdc42 inhibitor treatment and Cdc42 knockdown was assessed by measuring cleaved and total PARP levels in whole-cell extracts using immunoblotting analysis.

#### Puromycin-incorporation assay

Effect of Cdc42 knockdown on global protein translation was examined using puromycin-incorporation assay. Briefly, CCD-18Co cells and primary HIMF transfected with Cdc42-specific siRNA or scrambled control siRNA were treated with or without TGF-β1 for 48 hours and then exposed to puromycin (10 µg/ ml) for 1 hour. Subsequently, cells were lysed using RIPA buffer, and subjected to immunoblotting analysis using anti-puromycin antibody. Preincubation of cells with known protein synthesis inhibitor, cycloheximide (20 µM for 1 hour) followed by exposure to puromycin for 1 hour were used as positive control to determine the specificity of puromycin labeling of newly-synthesized polypeptides. A ‘no puromycin’ treatment group served as a negative control.

#### Immunofluorescence

For immunofluorescence, CCD-18Co cells and primary HIMF were plated on collagen-coated glass coverslips in 24-well plate and glass chamber slides, respectively. Following different treatments, cells were washed with PBS, fixed with 4% paraformaldehyde (PFA) for 15 minutes, and permeabilized in 0.5% Triton X-100 for 5 min at room temperature. Fixed cells were blocked with 1% BSA for 1 hour at room temperature followed by incubation with 1 hour (α-SMA, collagen I or fibronectin, dilution: 1:200) or overnight (p-SMAD2, dilution: 1:100) incubation. The samples were washed three times with the blocking buffer (1% BSA) followed by incubation with Alexa-Fluor-568–conjugated donkey anti-mouse or anti-rabbit secondary antibodies at 1:500 dilution in the blocking buffer for 1 hour. After secondary antibody incubation, cells were washed three times with the blocking buffer and mounted with ProLong™ Gold Antifade mounting medium with DNA stain DAPI (Thermo Fisher Scientific, P36941). For the actin cytoskeleton labeling, fixed/permeabilized cells were incubated for 1 hour with Alexa-Fluor-488 conjugated phalloidin. Images were captured using 40× oil immersion objectives (1.25 NA) on a laser scanning confocal microscope (DMi8 inverted microscope equipped with TCS SP8 confocal scanner (Leica Microsystems, Wentzler, Germany), using LAS-X acquisition software (Leica Microsystems, v3.5.5).The signal intensities of fluorescence images were measured using the Image J software (National Institute of Health, Bethesda, MD).

### Quantification and statistical analysis

Data are presented as the mean ± standard error of mean (SEM). The statistical analysis was performed by using two-tailed unpaired Student’s test to compare the results obtained with two experimental groups. Multiple groups were compared using one-way or two-way ANOVA, followed by Tukey or Holm-Šídák post-hoc test, as specified in the figure legends. Kruskal-Wallis test was used for multiple comparisons, followed by Dunn’s post-hoc test, as specified in the figure legend. All statistical analysis was performed using GraphPad Prism 11.0. Detailed statistical methods are provided in the figure legends. *p* values < 0.05 were considered statistically significant (represented as ^∗^*p* < 0.05, ^∗∗^*p* < 0.01, ^∗∗∗^*p* < 0.001, or not significant (ns)).

## Supplementary figure legends

**Supplementary figure 1.**
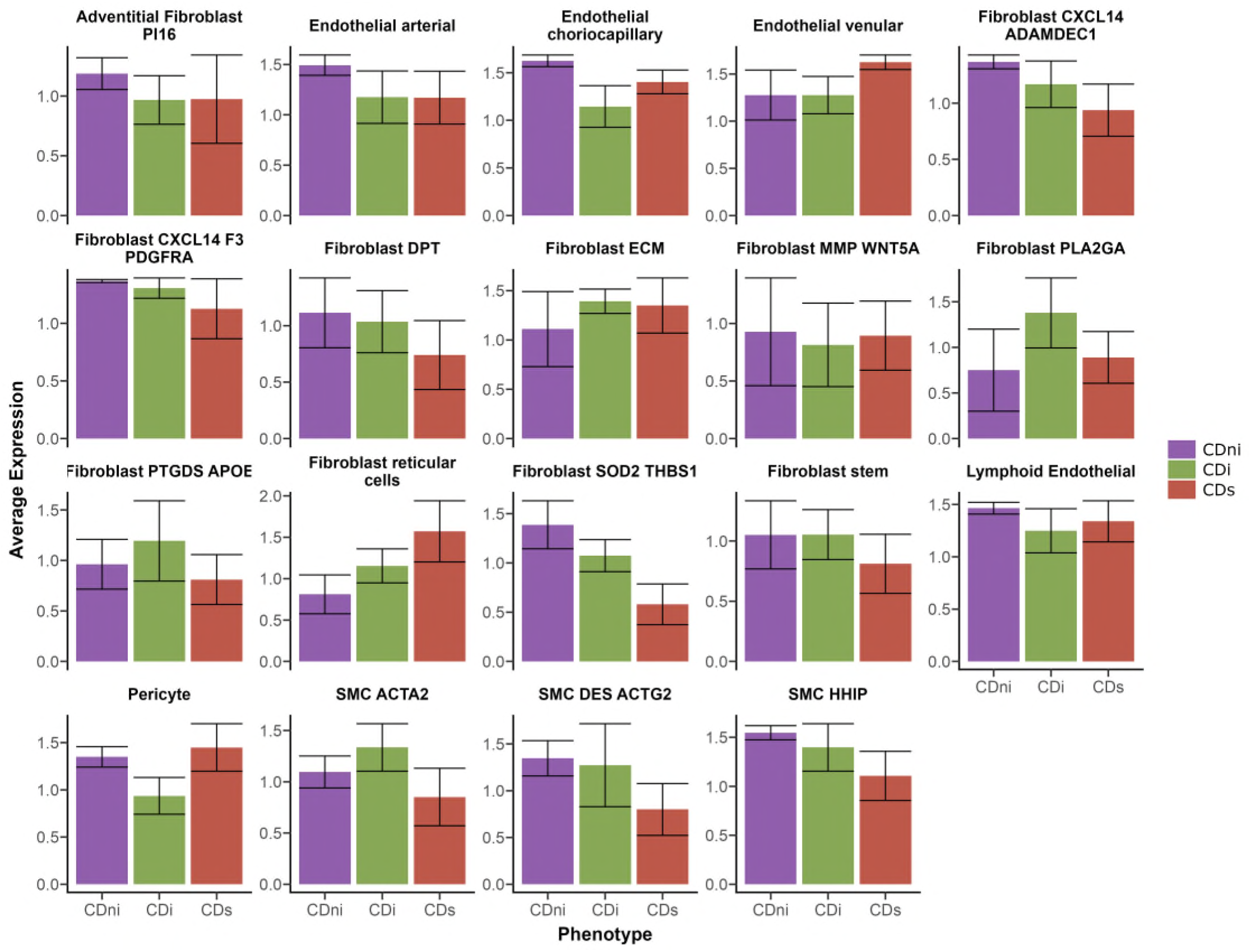
Cdc42 expression in stromal cell populations in fibrotic versus non-fibrotic tissues of Crohn’s disease (CD) patients. CDni - nonstrictured, noninflamed Crohn’s disease; CDi - nonstrictured but inflamed Crohn’s disease; CDs - strictured Crohn’s disease. Panels were generated from single cell RNA sequencing data described in our prior publication^37^.

**Supplementary figure 2.**
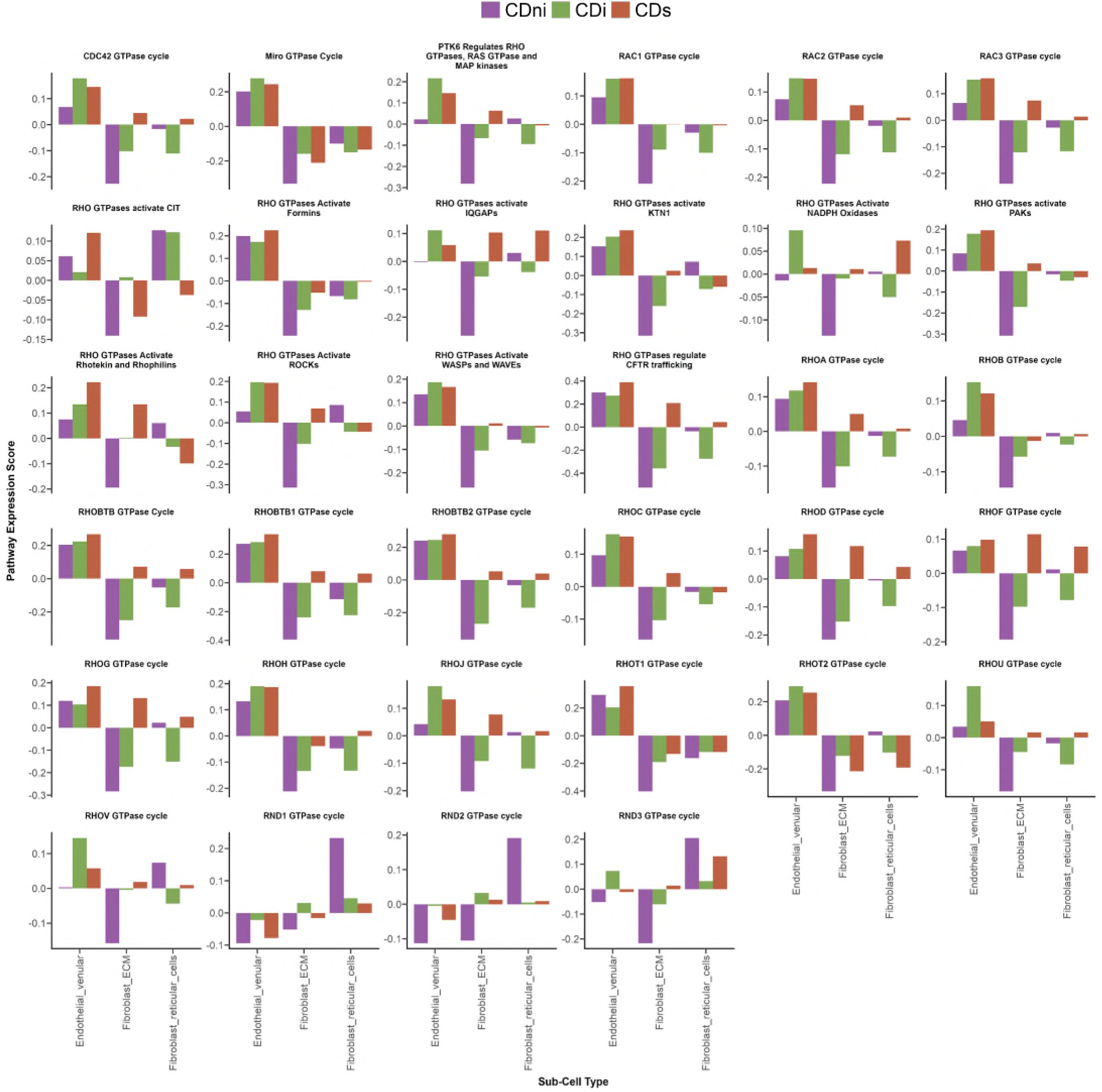
Pathway enrichment score of Cdc42-high stromal cells in strictured Crohn’s disease (CD) patients. CDni - nonstrictured, noninflamed Crohn’s disease; CDi - nonstrictured but inflamed Crohn’s disease; CDs - strictured Crohn’s disease. Panels were generated from single cell RNA sequencing data described in our prior publication^37^.

**Supplementary figure 3.**
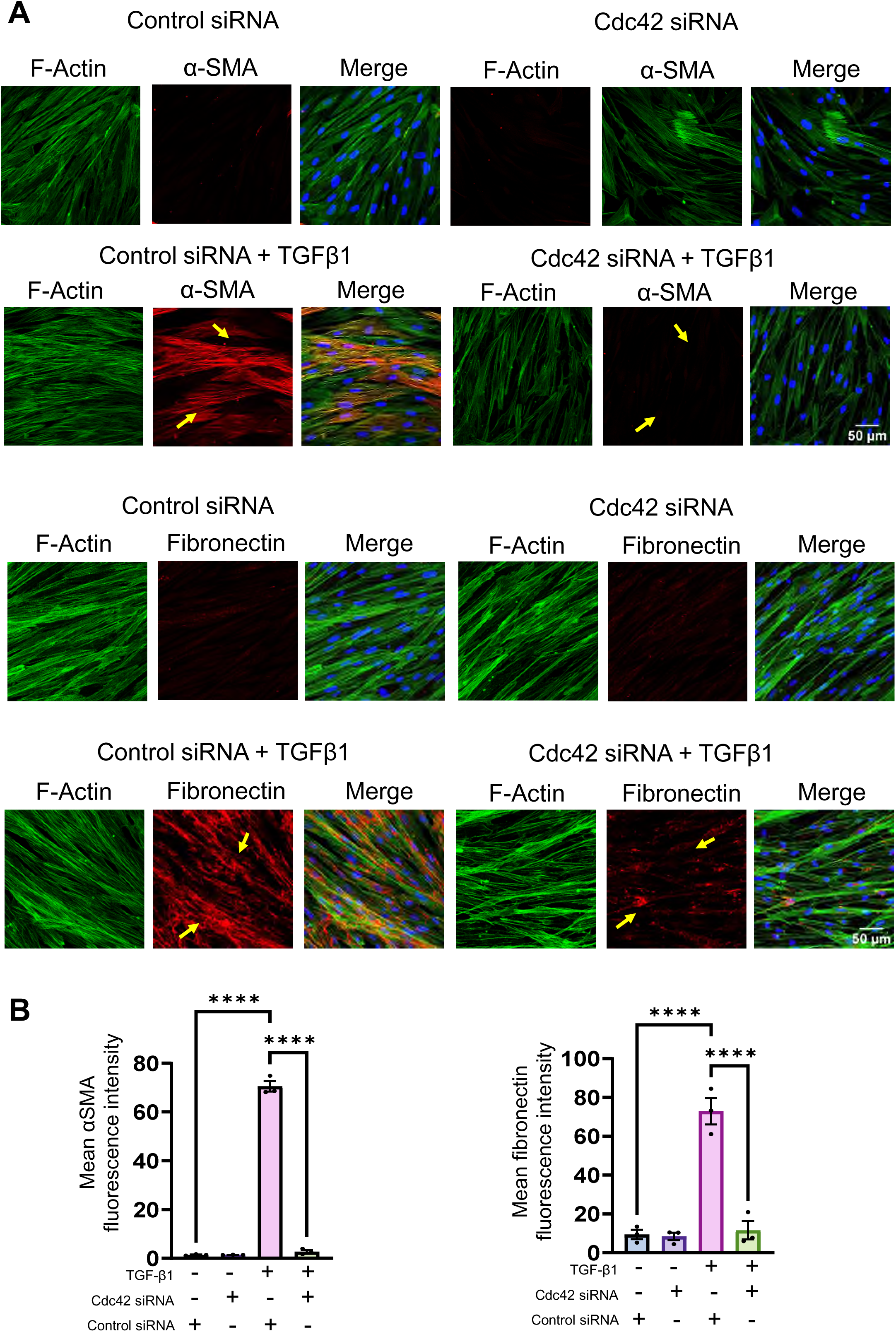
Cdc42 knockdown inhibits TGF-β1-induced activation of primary HIMF isolated from CD stricture. HIMF isolated from strictured CD patients were transfected with control or Cdc42-specific siRNA and stimulated with TGF-β1 for 48 hours. Immunofluorescence labeling and confocal microscopy was used to examine α-SMA and fibronectin expression in control and Cdc42 depleted cell under resting and TGF-β1-stimulated conditions. **(A)** Images shown are representative of three independent experiments. **(B)** Mean fluorescence intensity of images was measured using the Image J software. Arrows indicate αSMA and fibronectin induction in TGF-β1 treated CD primary HIMF transfected with scrambled control siRNA and inhibition of αSMA and fibronectin in HIMF stimulated with TGF-β1 transfected with Cdc42-specific siRNA. Data are presented as mean ± SEM (n=3). *p < 0.05; **p < 0.01; ***p < 0.001 by one-way ANOVA with Tukey’s multiple-comparison test. Scale bars, 50 µm.

**Supplementary figure 4.**
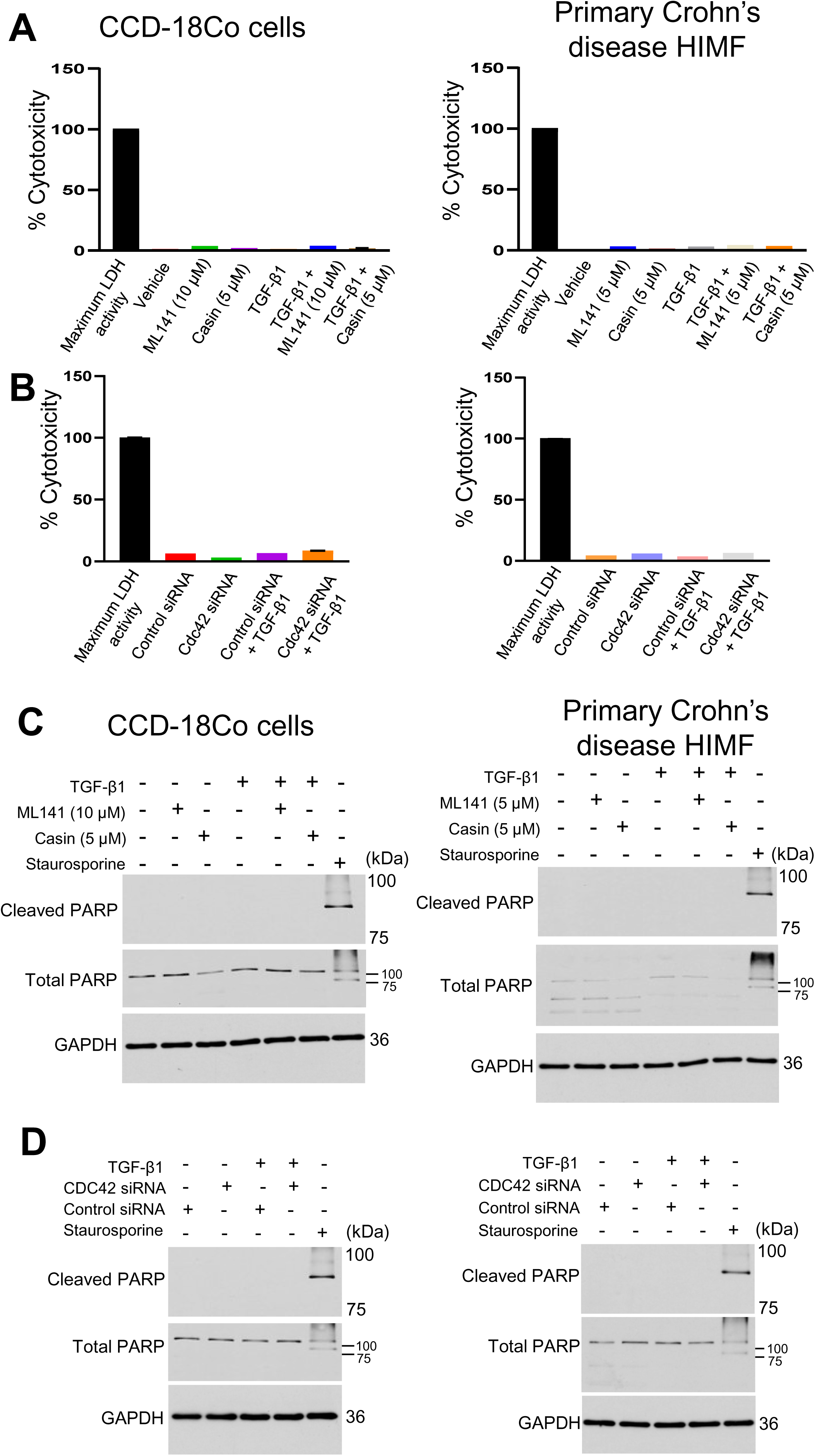
Cellular toxicity assessment of Cdc42 inhibitors and knockdown of Cdc42 in intestinal fibroblasts with or without TGF-β1 stimulation. (A and. **B)** Cytotoxicity was assessed by LDH release in CCD-18Co and primary HIMF either treated with ML141 or casin at indicated concentrations, or transfected with control or Cdc42-specific siRNA for 48 hours with or without TGF-β1 stimulation. Maximum LDH activity represents treating fibroblasts with lysis buffer for 1 hour. Data are presented as mean ± standard deviation (SD). (**C and D**) Cellular cytotoxicity following Cdc42 inhibitor treatment or Cdc42 knockdown was assessed by measuring levels of cleaved and total PARP in whole-cell lysates using immunoblotting. Cell treatment with a classical apoptosis inducer, staurosporine, served as a positive control for the PARP cleavage.

**Supplementary figure 5.**
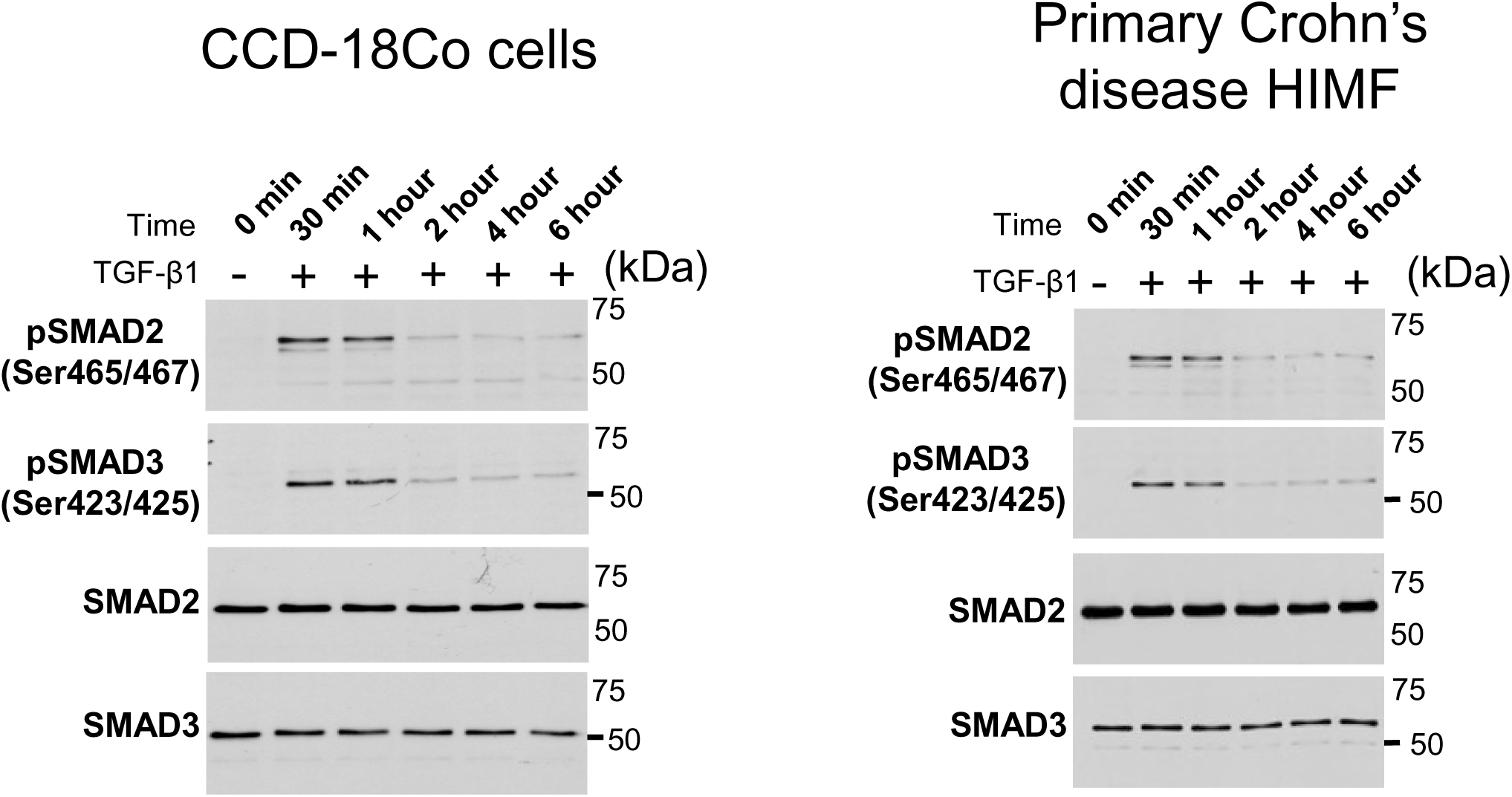
TGF-β1 induces early Smad activation in intestinal myofibroblasts. CCD-18Co cells and primary CD HIMF were stimulated with TGF-β1 for the indicated times. Immunoblotting analysis shows expression of total and active (phosphorylated) SMAD2 and SMAD3 in total cell lysates.

**Supplementary figure 6.**
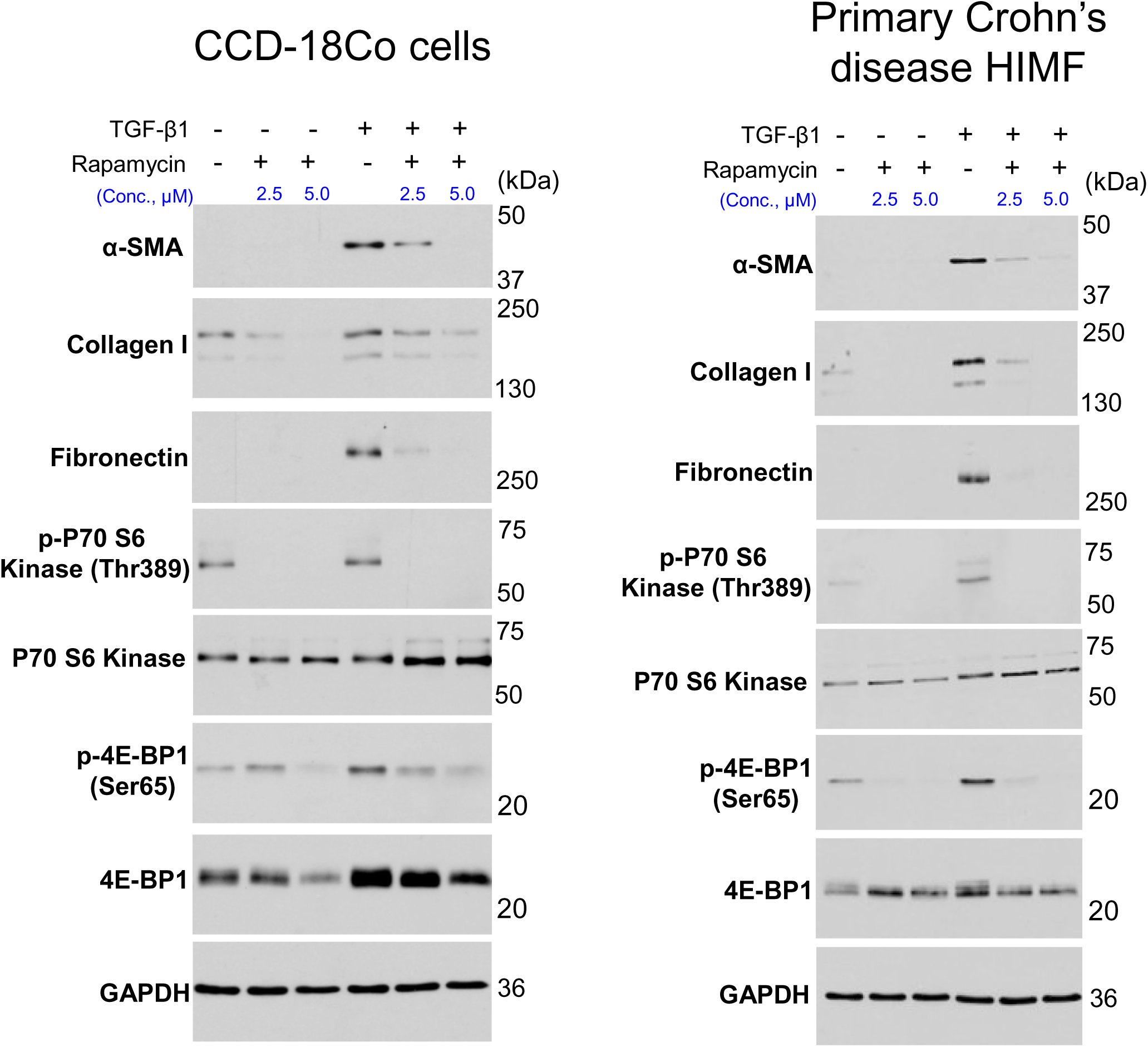
mTOR inhibition attenuates TGF-β1 induced activation of intestinal myofibroblasts. CCD-18Co cells and CD HIMF were stimulated with TGF-β1 for 48 hours in the presence or absence of increasing concentrations of rapamycin (0-5.0 µM). Immunoblotting analysis shows the expression of α-SMA, collagen I and fibronectin along with expression of total and activated downstream mTOR effectors (p70 S6 kinase and 4E-BP1) in total cell lysates.

**Supplementary figure 7.**
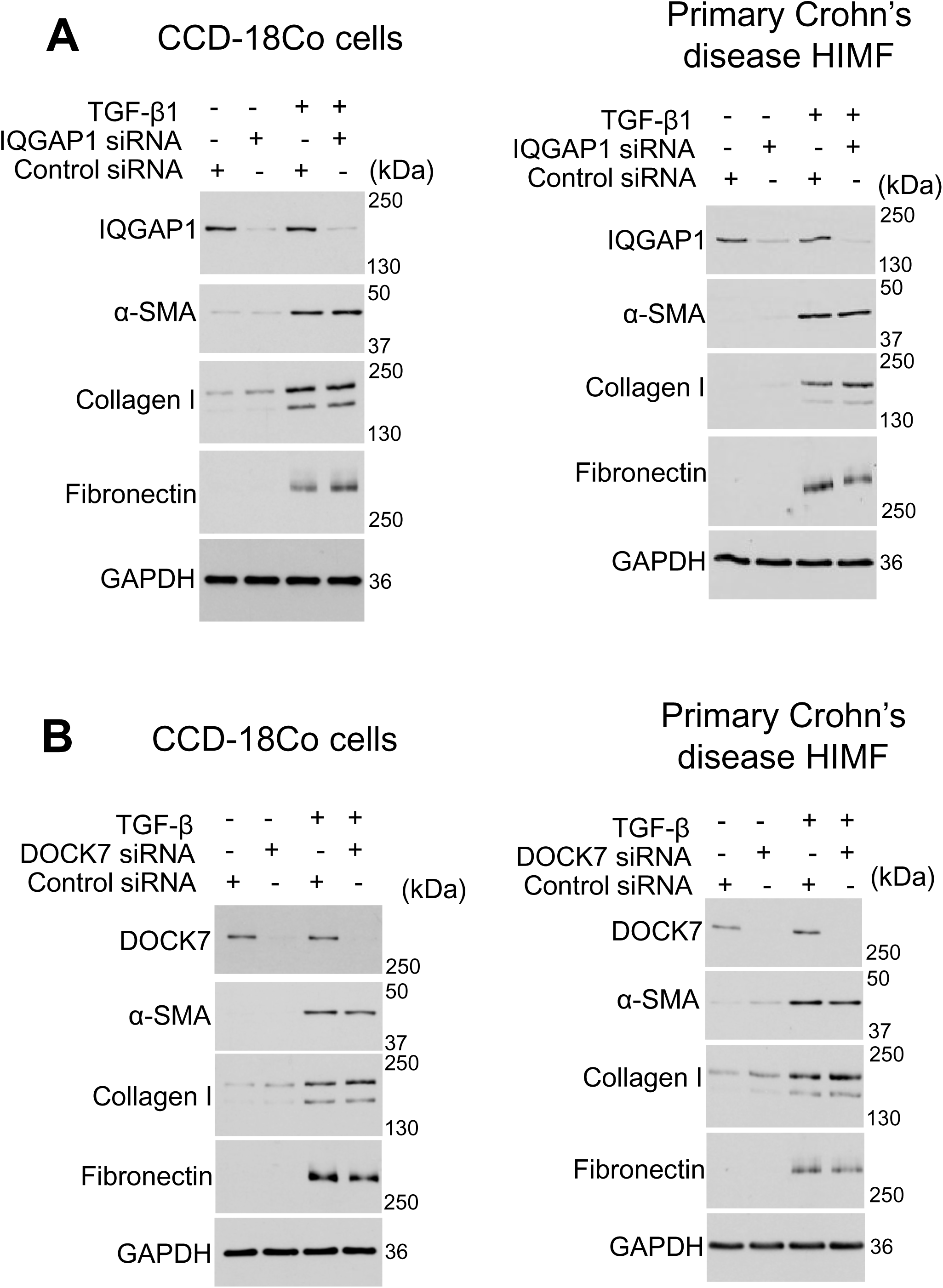
Loss of Cdc42 scaffolding proteins (IQGAP1 and DOCK7) does not affect TGF-β induced activation of intestinal myofibroblasts. CCD-18Co cells and primary CD HIMF were transfected with either control, IQGAP1 **(A)** or DOCK7 **(B)** siRNAs and stimulated with TGF-β1 for 48 hours. Immunoblotting analysis shows the efficiency of knockdown and expression of α-SMA, collagen I and fibronectin in resting and TGF-β1 stimulated control and IQGAP1 or DOCK7 depleted intestinal myofibroblasts.

